# Semaphorin3f as an intrinsic regulator of chamber-specific heart development

**DOI:** 10.1101/2021.05.19.444704

**Authors:** R Halabi, P.B. Cechmanek, C.L. Hehr, S. McFarlane

**Affiliations:** Graduate Program in Neuroscience, University of Calgary; Hotchkiss Brain Institute, Alberta Children’s Hospital Research Institute, Department of Cell Biology and Anatomy, University of Calgary. 3330 Hospital Dr., NW, Calgary, AB, Canada, T2N 4N1

## Abstract

During development a pool of precursors form a heart with atrial and ventricular chambers that exhibit distinct transcriptional and electrophysiological properties. Normal development of these chambers is essential for full term survival of the fetus, and deviations result in congenital heart defects. The large number of genes that may cause congenital heart defects when mutated, and the genetic variability and penetrance of the ensuing phenotypes, reveals a need to understand the molecular mechanisms that allow for the formation of chamber-specific cardiomyocyte differentiation. We find that in the developing zebrafish heart, mRNA for a secreted Semaphorin (Sema), Sema3fb, is expressed by all cardiomyocytes, whereas mRNA for its receptor Plexina3 (Plxna3) is expressed by ventricular cardiomyocytes. In *sema3fb* CRISPR zebrafish mutants, ventricular chamber development is impaired; the ventricles of mutants are smaller in size than their wild type siblings, apparently because of differences in cell size and not cell numbers, with ventricular cardiomyocytes failing to undergo normal developmental hypertrophy. Analysis of chamber differentiation indicates defects in chamber specific gene expression at the border between the ventricular and atrial chambers, with spillage of ventricular chamber genes into the atrium, and vice versa, and a failure to restrict *bmp4a* mRNA to the atrioventricular canal. The disrupted atrioventricular border region in mutants is accompanied by hypoplastic heart chambers and impaired cardiac function. These data suggest a model whereby cardiomyocytes secrete a Sema cue that, through spatially restricted expression of the receptor, signals in a ventricular chamber-specific manner to establish a distinct border between atrial and ventricular chambers that is important for functional development of the heart.

## Introduction

The heart is the first organ to form and become functional in an embryo, and congenital heart defects affect 9 per 1000 live births^1^. Collectively, congenital heart defects involve the irregular formation of heart chambers and/or valves. In vertebrates, chamber morphogenesis follows the twisting, expansion and septation of the linear heart tube into a two- (teleost), three- (amphibian) or four-(mammalian/avian) chambered heart^2^. These cardiac chambers, the atrium and ventricle, differ in their biochemical properties, electrophysiological and contractile capacities, as well as in their transcriptional profiles^3, 4^. What maintains the segregation of the cell populations in the two discrete chambers as the embryonic heart undergoes morphogenesis is unknown.

During development, cells within tissues acquire unique identities, and need to be segregated from one another within subdivisions of the tissue^5, 6^. This is true for example of the developing hindbrain, where the initial expression of genes within adjacent rhombomeres is initially imprecise, and is sharpened as embryogenesis proceeds, producing domains with straight borders^6^. Key in this process are interactions between neighbouring hindbrain cells that involve contact-mediated repellent signaling via Ephrin ligands and Eph receptors. Whether a similar process of refinement of the border occurs between ventricular and atrial chambers of the heart is unknown. Chamber-specific development depends on extrinsic factors, which include Nodal, Notch, Bone morphogenetic protein (Bmp), Fibroblast growth factor (Fgf) and retinoic acid^7–9^. These signals then pattern the early heart tube to establish secondary localized signaling that spatially restricts the expression of differentiation genes to the specific chambers and valves. Intrinsic regulators, such as transcription factors, are also necessary for chamber-specific cardiomyocyte development. For example, ventricle specific expression of Irx4 is necessary in chick for the expression of ventricular Myosin Heavy Chain I (MHC1) and suppression of atrial MHC^10^. Further, in mouse PITX2 within the left atrium inhibits the expression of *Shox2* and the specification of the pacemaker cells of the sinoatrial node^11^. An unanswered question is whether active mechanisms are required to maintain segregation of the two populations of cardiomyocytes once established, especially in light of the disruptive forces of cardiac chamber morphogenesis and heart looping.

One group of molecules known to act as repellents for developing cells of the nervous and cardiovascular systems are the secreted class 3 Semaphorins (Sema3s), which are used as guidance cues for vessels, axons and neural crest cells^12–15^. Sema3 signaling is mediated through the canonical Plexin (Plxn) receptors and their coreceptors, the Neuropilins (Nrp)^16^. Sema3s are implicated in murine cardiovascular development via their regulation of the directed movement of associated cells; SEMA3C^17^, SEMA3D^18^, SEMA3E^19^, and SEMA3G^20^ pattern embryonic vessels, SEMA3A regulates sympathetic innervation of the heart^21^, and SEMA3C promotes neural crest cell migration into the heart^17^. Whether Sema3s play roles in heart formation via the direct regulation of cardiomyocyte development is unknown.

Here, we take advantage of the zebrafish model as a powerful tool to study the development of the heart; the embryos do not require a functional cardiovascular system during embryogenesis^22^, which allows for the extended characterization of manipulations that are normally fatal. We find that mRNA for the secreted Sema3, Sema3fb, is expressed early in cardiomyocytes during chamber differentiation, while *nrp2b* and *plxna3* receptor mRNAs are expressed by ventricular cardiomyocytes, suggesting spatially restricted Sema3fb signaling within the developing myocardium. To investigate a role for ventricular cardiomyocyte Sema3fb signaling, we generated CRISPR/Cas9 zebrafish *sema3fb* mutants. With either loss of Sema3fb, or signaling through its Plxna3 receptor via morpholino-oligonucleotide knockdown, chamber specific gene expression is either lost from or expands beyond the border between the atrium and ventricle. Accompanying the loss of segregated gene expression between the cardiac chambers is a decrease in the size of both chambers, and significantly impaired cardiac output and function of the larval heart. We propose a model whereby Sema3fb signaling ensures segregation of atrial and ventricular cardiomyocyte chambers, a feature that appears critical for chamber development and heart function.

## Methods

### Zebrafish strains and maintenance

Zebrafish (Danio rerio) were maintained according to standard procedure on a 14-h light/10-h dark cycle at 28°C. Embryos were obtained by natural spawning, raised in E3 medium supplemented with 0.25 mg/L methylene blue, and staged by convention^23, 24^. To permit exact developmental staging, with all embryos synced, adult zebrafish spawned for 10 minutes only before embryo collection. Dishes were screened at 6 hours post fertilization (hpf) to remove embryos that showed delayed embryogenesis or unfertilized eggs. We used *sema3fb^ca305^* and *sema3fb^ca306^* genetic mutants. These lines were maintained as heterozygotes for all initial experiments, and thereafter maintained as homozygotes and wild type (WT) siblings for future embryo collection. The *Tg(-6.5kdrl:mCherry)^ci5^* line was used as an outcross for *sema3fb^ca305^* mutants to label endothelial cells^25^. Sex was not a variable in these studies as zebrafish cannot be sexed as embryos. The University of Calgary Animal Care Committee approved all procedures (#AC19-0114).

### Establishment of *sema3fb* mutant lines

To generate genetic knock-outs by using the CRISPR/Cas9 system a sgRNA (5’ - GAAGGACAAGAAGACCCGCG) targeting *sema3fb* exon 2 was selected following CHOPCHOP query^26^ and analysis of secondary structure using Vienna RNAfold Prediction (rna.tbi.univie.ac.at). sgRNA template generation and transcription was carried out in accordance to the protocol described previously^27^. 1-cell stage embryos were injected with a 1 nl mix of approximately 56-60 pg sgRNA and 190 pg *cas9* mRNA (Addgene plasmid #47322). Mosaic embryos were raised to adulthood and crossed with Tupel Long fin (TL) fish to identify founders. To genotype, caudal fin clippings were collected from tricaine methanesulfonate (MS-222; 160mg/L) anesthetized adults. Genomic DNA extraction was performed^28^ and amplified by PCR using primers (Forward: 5’- ATTGCCCCACAAAATAACATTC; Reverse: 5’ - GTCTACTCTGTGAATTTCCCGC) around the expected mutation site. Amplicons were sequenced at University of Calgary DNA Core facility for definitive genotype confirmation.

### RNA extraction and quantitative real time PCR

Total RNA from embryos was prepared using the TRIzol reagent (Invitrogen) extraction method^29^. Briefly, 5 embryos or embryonic hearts at specific developmental ages were collected in Trizol and homogenized using a 26½ gauge syringe. First strand cDNA was made using the Superscript II RT-PCR (Invitrogen, 11904-018) protocol with 100 ng of total RNA for conversion. Each RT-qPCR reaction had a 10 µl final volume containing 0.5 µl of cDNA, 500 nM of each primer and 5 µl of SYBR Green QuantiFast RT-qPCR master mix (Qiagen). Primers used were: *sema3fb* (F: 5’- AGTGACGCATATGGCTCTGC; R: - 5’- AGGAAGCCTCTTCTGCGAGG); *bmp4a* (F: 5’ – GACCCGTTTTACCGTCTTCA; R: 5’- TTTGTCGAGAGGTGATGCAG) (Morris et al., 2011); *tbx5a* (F: 5’- AACCATCTGGATCCCTTCG; R: 5’- TGTTTTCATCCGCCTTGAC). An Applied Biosystems QuantiStudio 6 Real-Time PCR system was used, and gene expression was normalized relative to reference gene *ß-actin* (F: 5’-GCAGAAGGAGATCACATCCCTGGC; R: 5’- CATTGCCGTCACCTTCACCGTTC). Reactions were performed in three technical replicates, and all results made from 3-4 independent biological replicates.

### *in situ* hybridization

Digoxigenin RNA probes were synthesized as described previously^30^. Probes were generated from plasmid templates or via PCR products containing an RNA polymerase binding sequence on the reverse primer. Probes used included: *bmp4a* (5’- CTGCCAGGACCACGTAACAT; 5’- GAAATTAATACGACTCACTATAGGGTGGCGCCTTTAACACCTCAT); *crip2* (5’-GCCCCAGATGCAGCAAGAAG; 5’-TCATTTAGGTGACACTATAGACCAGCACCAGTCACAAACACC); *irx1a* (5’-GAGAACAAGGTGACCTGGGG; 5’-GAAATTAATACGACTCACTATAGGGTGAAGAGGACGAAACGACGA); *ltbp3a* (5’ACTCACCTTTAGTGCAGCCC; 5’-GAAATTAATACGACTCACTATAGGGCTTCAAATGGGCGCAAACCC); *myh6* (5’-GGAGTACGTGAAGGGGCAAA; 5’-GAAATTAATACGACTCACTATAGGGGCTCGTCCCGAAATGAATGC); *myh7* (5’-GCAACTTGGTGAGGGAGGAA; 5’-GAAATTAATACGACTCACTATAGGGAGCAAGCTTACGGCCTCTTT); *nrp2b* (5’-TCCGGGATGGGAACTCAGAT; 5’- GCATTTAGGTGACACTATAGAATCGGCCGTCATAGAAGCTG); *plxna3* (pCRII, linearized with EcoRV); *sema3fb* (pCR4, linearized with NotI); *tbx5a* (pCRII, linearized with EcoRV). Whole mount *in situ* hybridization was performed as described previously with some minor modifications^30^. We omitted steps with gradient changes in hybridization buffer, and the 2xSSC step was carried out at 70°C and 0.2 x SSC at 37°C. Embryos were incubated directly in anti-DIG-AP, Fab fragment antibody (Roche) solution in blocking buffer at room temperature for 2 hours with no blocking step. NBT/BCIP (Roche) was used at a 2.5/3.5µl/ml ratio to stain embryos. Stained embryos were fixed in 4% paraformaldehyde (PFA), and washed in PBS with 0.1% Triton X-100 (PBST). When directly comparing *in situ* hybridization signals between genotypes, all embryos were processed in the same tube.

### Immunolabeling and Imaging

Embryos were fixed overnight in 4% PFA in PBS and processed for whole mount immunostaining. Briefly, samples were blocked in PBST with 10% bovine serum albumin and 2% normal sheep serum for 30 minutes and transferred to primary antibody solution in 1/10 blocking buffer overnight at 4°C. Samples were then washed with PBST and incubated in secondary antibody (Alexa Flour 488/555 rabbit/mouse) supplemented with 5 mg/ml Hoechst in PBST for 45 minutes. Embryos were fixed and imaged by confocal microscopy. The following primary antibodies were used: MF20 (1:500, Developmental Studies Hybridoma Bank), pHH3 (1:500, Millipore), zn8 (DM-GRASP; 1:500, Developmental Studies Hybridoma Bank). ApopTag Peroxidase In Situ Apoptosis Detection kit (Millipore) was used to detect apoptotic cells in whole mount embryos, with overnight diaminobenzidine staining. For confocal microscopy, embryos were imaged with a 10 or 20x objective on a Zeiss LSM 700 microscope after mounting in 0.8-1% low melting point agarose (Invitrogen) on a glass bottom dish. Optical slices were taken at intervals from 1-5 µm and subjected to 2 times averaging. Z-stacks were processed in Zen Blue as maximal projections and compiled using Adobe Photoshop 2020.

### Heart Function and Morphology Analysis

Embryos (72 hpf) were anaesthetized with minimal MS-222 and positioned on their right side laterally on a glass slide in a water droplet. Video images were taken 30 frames/second for 15 secs using a Leica DM5500 B microscope equipped with a Leica DFC365 FX. Heart rate was determined in a 15 sec window. Measurements^31^ were made from frame stills during systole and diastole across three heart beats using ImageJ. Briefly, the following equations were used^2^: Ventricle volume = (0.523) (Ventricle Width) (Ventricle Length); Fractional Shortening= (100) (Ventricle Width Diastole-Ventricle Width Systole)/ Ventricle Width Diastole); Stroke Volume = Ventricle Volume Diastole – Ventricle Volume Systole. Ejection Fraction = (100) (Stroke Volume/Ventricle Volume Diastole). Cardiac Output = (Heart Rate) (Stroke Volume).

### Hematoxylin and Eosin (H&E) staining

H&E staining of JB-4 plastic sections was carried out as described previously^32^ with the following modifications; Acid and base washes were omitted, Harris Modified Hematoxylin Solution (Sigma) and Eosin Y (EMD) were used, and regular tap water replaced prescribed substitute. Slides were air dried for 15 mins^33^ and coverslipped using Permount (Fischer Scientific). Myocardial ventricle wall thickness for each embryo was the average of measurements made at two points along the anterior-posterior axis from 1-2 sections stained by H&E.

### Cardiomyocyte measurements

Morphometries (area and perimeter) of confocal imaged, Zn8-immunolabelled cardiomyocytes were measured in ImageJ at 48 hpf. Cells measured had clearly visible outlines. Of note, only cells central to the outer curvature of the ventral ventricle were assessed. Circularity was used to distinguish cellular morphologies in which 1 represents a perfect circle. This was calculated by the following formula: Circularity = 4 πArea/Perimeter^2^. Measurements were averaged from 8-12 cells per embryo. Additionally, projected confocal MF-20-immunolabeled images were analyzed using ‘RGB Measure’ tool on ImageJ across the entirety of the heart, values averaged for all samples, and represented as an intensity plot across distance.

### Sema3f Overexpression and Knockdown

For heart specific overexpression, *sema3fa* was amplified from Image Clone 9038816 using primers that incorporate *attb1/b2* recombination sites (5’- GGGGACAAGTTTGTACAAAAAAGCAGGCTTCACCATGCAGGGAGCCGGGACT TTGGTG-3’ and 5’- GGGGACCACTTTGTACAAGAAAGCTGGGTTTGTCTCAGCCATGCTGCTCTGCTC-3’) and inserted into pDONR221 to create a *pME-sema3fa* vector. Three way Tol2 gateway cloning^34^ was used to insert *sema3fa* downstream of the cardiomyocyte *myl7* promoter^35^ giving *myl7:sema3fa:p3E-MTpA* in Tol2 with *pDest:mKate*^36^ as the transgenesis marker. One cell-stage zebrafish embryos were injected with a solution consisting of 12.5-25 ng/µl plasmid with or without 25 ng/µl *transposase* mRNA. At 48 hpf the embryos were checked for mKate eye expression and fixed for *irx1a* and *bmp4a in situ* hybridization as well as anti-myc immunohistochemistry. For knockdown of *sema3fb* we injected either 1 or 3 ng/ml of an antisense morpholino oligonucleotide that targets the ATG start site of *sema3fb* (<u>CAT</u>AGACTGTCCAAGAGCATGGTGC, Gene Tools LLC, Corvalis, OR, USA) into 1-cell zebrafish embryos. For *plxna3* and *nrp2b* knockdown we used previously characterized MOs^37^. At 48 hpf, embryos were scored for edema and fixed for *in situ* hybridization analyses.

### Statistics

All data sets for qualitative scoring or quantitation were analyzed in a blinded fashion. Sample size calculations were not performed because we characterized mutants, and thus had no prediction as to the likely size of effects to use in power calculations. Results are expressed as mean ± standard error the mean (SEM) unless otherwise indicated. All statistical analysis was performed using Prism 8 software (Graph Pad). An unpaired, non-parametric Mann Whitney U test was used for comparisons of two samples, and a One-Way ANOVA with a Kruskal-Wallis test allowed for comparisons between multiple samples.

## Results

### *sema3fb* is expressed by the cardiomyocyte progenitor cells of the developing heart

To elucidate the spatial and temporal expression of *sema3fb* in the zebrafish heart we performed whole mount RNA *in situ* hybridization (ISH) on embryos from 10-72 hours post fertilization (hpf). In zebrafish, cardiac progenitors of the atrial and ventricular chambers are bilaterally specified by 5 hpf^38^. The progenitors then condense at the midline and undergo a series of movements to produce a linear tube of differentiating myocardium that surrounds the underlying endocardium^38, 39^. This tube starts pumping between 24-26 hpf^40^. By 48 hpf, the two-chambered heart is discernable by the constriction of the atrioventricular canal^38, 41^. *sema3fb* transcript was first detected at 10-18 hpf in a bilateral fashion in the presumptive cardiac progenitor fields, as revealed by the cardiomyocyte progenitor marker *hand2* (Suppl Fig. 1B). By 24 hpf, *sema3fb* was expressed by cardiomyocyte progenitors through the entire linear heart tube (Figure 1A), as evidenced by ISH signal similar to that seen with the pan-cardiomyocyte marker *cardiac troponin T type 2a* (*tnnt2a*) (Figure 1B)^42^. A thin transverse plastic section of *sema3fb* whole mount ISH on a *Tg(flk:EGFP)* background, where GFP labels the endocardial cells that line the ventricles, revealed that *sema3fb* was expressed not by GFP positive cells but the cardiomyocytes of both atrial and ventricular chambers (Figure 1C-E). *sema3fb* remained expressed in both the ventricle and the atrium over the period that the heart undergoes rightward looping and chamber morphogenesis (24-48 hpf) (Figure 1F). Weak expression persisted throughout the ventricular and atrial myocardium of the 72 hpf heart (data not shown). These data indicate that mRNA for a secreted Sema is expressed by cardiomyocytes at the time they undergo differentiation, and agree with RNAseq analysis of FACs-isolated zebrafish cardiomyocytes, where *sema3fb* mRNA is enriched in the heart over the rest of the embryo^43^. The presence of *sema3fb* in cardiomyocytes suggests a role for Sema3fb in heart development in a novel heart-intrinsic manner.

**Figure 1:**
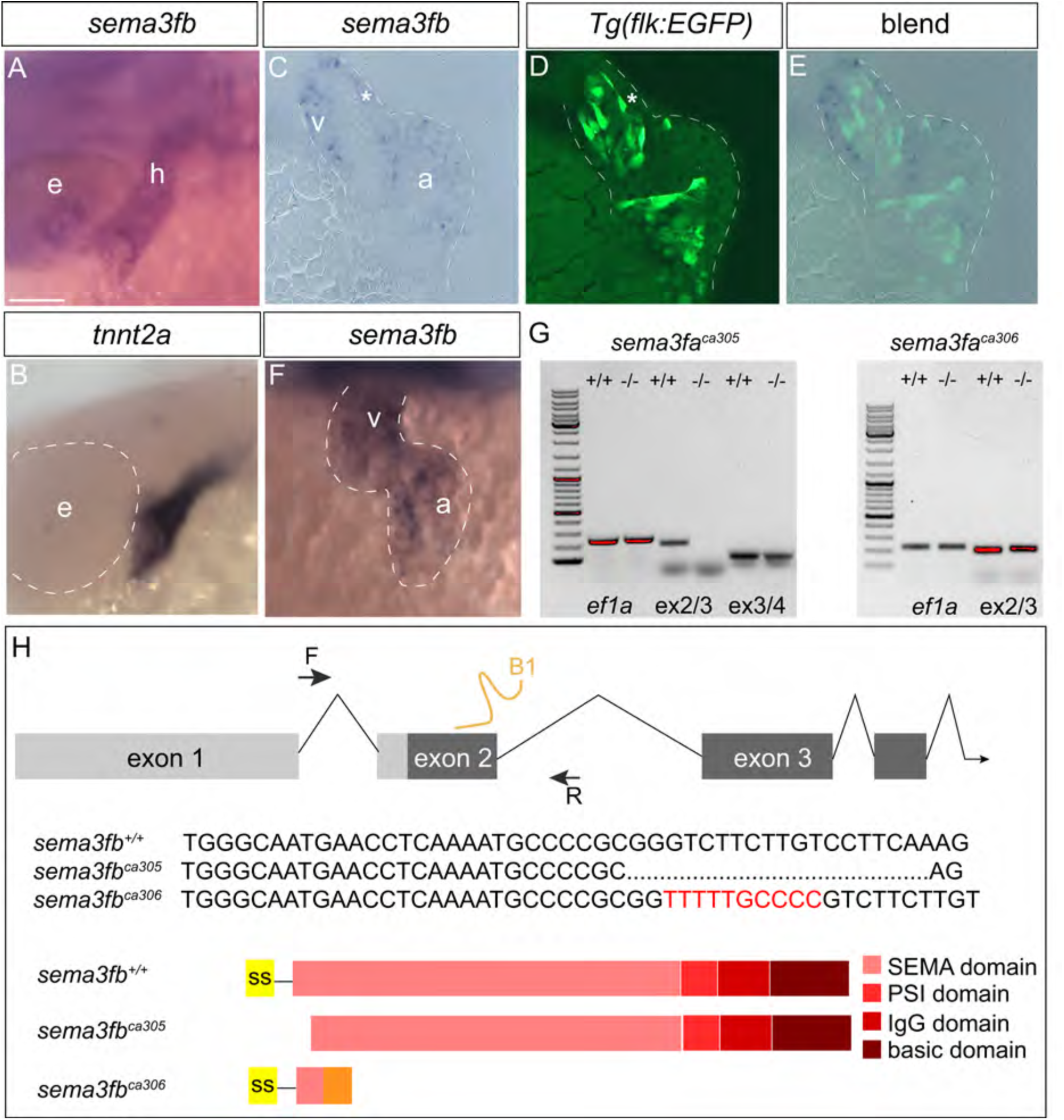
*sema3fb* is expressed by cardiomyocytes of the developing heart. **A-F)** Whole mount ISH of 28 hpf (A,B) and 48 hpf (F) zebrafish hearts labeled with antisense riboprobes for *sema3fb* (A,F) and the cardiomyocyte marker *tnnt2a* (B). Transverse section through the heart at 36 hpf reveals *sema3fb* ISH signal present in both chambers of the heart (C) in cardiomyocytes (asterisk) and not the endocardium as labeled by GFP (D) in a *Tg(flk:EGFP)* background. Blend of C and D shown in E. **G)** RT-PCR analysis indicates that exon 2 is skipped in the *sema3fb^ca305^* allele; no PCR product is generated with a forward primer in exon 2, but PCR product is generated with an exon 3-exon 4 primer combination. In contrast, exon 2 is present in the *sema3fb^ca306^* allele, as demonstrated by a PCR product generated with an exon 2-exon 3 primer combination. **H)** Chromosomal overview of the *sema3fb* locus targeted by CRISPR/Cas9 mutagenesis to exon 2 (guide RNA: B1) and primers used to identify the mutation. *sema3fb^ca305^* mutants have a 19 bp deletion (dots) and *sema3fb^ca306^* mutants have a 10 bp insertion (red sequence). Schematic representation of WT and mutant proteins. In the *sema3fb^ca305^* allele, exon 1 is spliced in frame with exon 3, but, with the loss of exon 2, is missing 62 amino acids (aa) at the beginning of a protein that would be translated; sequence analysis indicates that both the signal sequence (ss) and the first 14 aa of the SEMA domain are missing. In the *sema3fb^ca306^* allele, the 10 bp insertion introduces a frame shift and a premature stop codon to generate a predicted truncated 58 aa protein. F, forward primer; R, reverse primer; UTR, untranslated region. A, anterior; a, atrium; e, eye; h, heart; P, posterior; v, ventricle. Scale bar: 50 µm.

To assess the importance of Sema3fb for heart development and function we used CRISPR/Cas9 gene editing technology^27^ to generate *sema3fb* genetic mutants. A specific guide RNA (sgRNA) targeting exon 2 (Figure 1H) was co-injected with *cas9* mRNA at the one cell stage and a single founder was recovered that housed two allelic variants; a 19 base pair (bp) deletion and a 10 bp insertion. Both lines were propagated from this founder to generate stable homozygous viable generations, alongside wild type (WT) siblings. RT-PCR analysis of mRNA isolated from 24 hpf embryos indicated that the 19 bp deletion mutant (hereafter *sema3fb^ca305^*) lacked exon 2 that contains the ATG start site, presumably because the deletion at the end of exon 2 disrupted a key splice site (Figure 1G). Indeed, sequencing of the PCR product generated with primers in exon 1 and exon 3 indicated that exon 1 was spliced in frame with exon 3 to gives an mRNA that if translated would produce a protein missing the first 62 amino acids (aa), including the signal sequence (amino acids 1-23) required for protein secretion and the first 14 aa of the SEMA domain that is necessary to elicit intracellular signaling^44^ (Figure 1H). Software could not identify another signal sequence in the predicted translated protein, thus in the *sema3fb^ca305^* allele the C-terminal truncated Sema3fb would not be secreted from cardiomyocytes. RT-PCR and sequencing analysis indicated that exon 2 was included in the *sema3fb* mRNA product in the 10 bp insertion allele (hereafter *sema3fb^ca306^*) (Figure 1G). The *sema3fb^ca306^* allele is predicted to produce a protein that is 58 aa in length and is prematurely truncated within the SEMA domain (Figure 1H). Of note, a commercial antibody against mouse SEMA3F failed to distinguish between multiple Sema3s in zebrafish (data not shown). Importantly, while injection of an antisense morpholino oligonucleotide (MO) against the ATG start site of *sema3fb* produced a similar heart edema when injected at the one cell stage in WT embryos as seen in the *sema3fb^ca305^* allele, there was no greater incidence of severe edema seen when the MO was injected into the *sema3fb^ca305^* allele (Suppl Figure 1A). The MO data argue strongly for the specificity of the heart phenotype in the *sema3fb* mutants being due to the loss of Sema3fb. Of note, *sema3fb* mRNA was not detected in the 48 hpf WT and *sema3fb* mutant heart (Suppl Figure 1C), arguing that there is no compensation for the loss of Sema3fb by upregulation of Sema3fa expression.

### Embryos that lack Sema3fb exhibit cardiac edema

To examine potential heart defects with the loss of Sema3fb, heterozygous adults were incrossed, and progeny raised to homozygosity and incrossed with their respective genotypes (*sema3fb^+/+^, sema3fb^ca305^, sema3fb^306^*). At 72 hpf, embryos were indistinguishable by gross size, but homozygous embryos exhibited edema (Figure 2A) that presented in both allelic variants at 48 hpf (8.3% *sema3fb*^+/+^, 91.7% *sema3fb^ca305^*, 76.7% *sema3fb^ca306^*, N=3, n=60 each genotype) and 72 hpf (5.0% *sema3fb*^+/+^, 91.7% *sema3fb^ca305^*, 41.7% *sema3fb^ca306^*, N=3, n=60 each genotype) (Figure 2B). Most mutant embryos exhibited edema, though the severity of the defect varied. Generally, embryos with edema developed to adulthood, though a small proportion (∼10%) of homozygous embryos did not survive past 5-7 days post fertilization (dpf), and exhibited severe cardiac edema (Figure 2C,D), a hypoplastic ventricle and a linear arrangement of the heart tube (Figure 2C’). All analyses were carried out on embryos in which the pericardial edema was of average extent, with lethal phenotypes excluded so as not to confound data interpretation. Of note, heterozygous *sema3fb^ca305^* and *sema3fb^ca306^* embryos also exhibited a spectrum of edema, ranging from no edema to severe edema, suggesting that genetic dosage influences the phenotype (data not shown). Additionally, as the phenotypes between the two allelic variants were highly comparable, we focused on the *sema3fb^ca305^* mutant in our analyses, and confirmed key findings in the *sema3fb^ca306^* allele. From hematoxylin and eosin (H&E) sections we found that the thickness of the ventricle myocardium at 72 hpf was reduced significantly (6.3 ± 0.5 µm, standard error of the mean (s.e.m.) here and throughout the paper, unless otherwise indicated; Mann Whitney U, p=0.024, N=2, n=6) in *sema3fb^ca305^* as compared to WT (9.4 ± 0.7 µm; N=2, n=3) hearts (Figure 2E), suggesting that in addition to edema, myocardial development was impaired.

**Figure 2.**
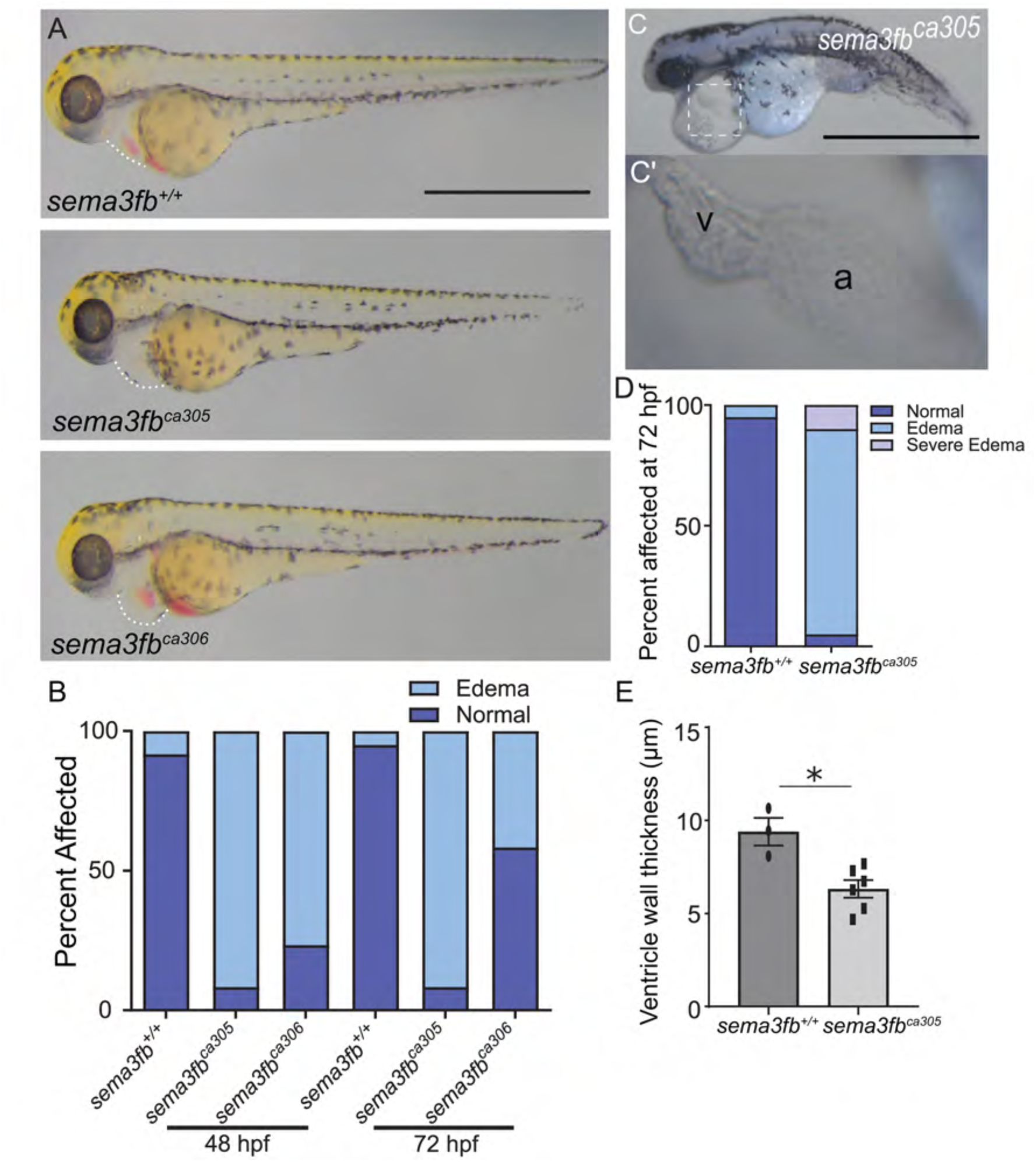
*sema3fb* mutants have cardiac edema. **A)** Representative brightfield images of 72 hpf *sema3fa*^+/+^, *sema3fb^ca305^* and *sema3fb^ca306^* embryos. Dotted line outlines the heart region. **B)** The percent of embryos of the different genotypes that present with cardiac edema at 48 hpf (N=3, n=60 for each genotype) and 72 hpf (N=3, n=60 for each genotype). **C)** Representative brightfield images showing the severe mutant phenotype that displays major cardiac edema and failure to thrive (C), and the abnormal linear arrangement of the heart (C’, inset in C). **D)** Quantitation of the edema phenotype at 72 hpf (N=3, n=180 per genotype). **E)** Quantitation reveals that the average ventricle wall width (µm) in *sema3fb^ca305^* embryos is significantly thinner (*p=0.024) than that of WT hearts. Error bars represent SEM. Statistics represent the Mann-Whitney U test. Scale bar in A and C is 1 mm.

### *sema3fb* mutants have impaired cardiac function

By 72 hpf in zebrafish, cardiac function is considered rhythmic and the circulatory system relatively mature^45^. To determine whether the edema in *sema3fb^ca305^* mutants resulted from impaired cardiac function, 72 hpf hearts were imaged by live video microscopy, and measurements of the ventricle were made at end-diastole and end-systole (Figure 3A-B). Ventricle volume was reduced significantly during diastole (0.50 ± 0.03 nL; p=0.0027) and systole (0.32 ± 0.02 nL; Mann Whitney U, p=0.0063) in *sema3fb^ca305^* mutants (N=3, n=20) as compared to WT siblings (diastole, 0.68 ± 0.05 nL; systole, 0.39 ± 0.02 nL; N=3, n=12) (Figure 3C-D). Additionally, mutants had a small but significantly slowed heart rate (WT 151.7 ± 1.4 beats/min, N=3, n=12; *sema3fb^ca305^* 143.0 ± 1.5 beats/min, N=3, n=20; Mann Whitney U, p=0.0008) (Figure 3E). Cardiac output (WT 44.5 ± 7.4 nL/min, N=3, n=12; *sema3fb^ca305^* 25.7 ± 2.6 nL/min; Mann Whitney U, p=0.021) and stroke volume (WT 0.29 ± 0.05 nL, N=3, n=12; *sema3fb^ca305^* 0.18 ± 0.02 nL, N=3, n=21; Mann Whitney U, p=0.04) (Figure 3F-G) were also reduced significantly in mutants as compared to WT, while ejection fraction (WT 40.5 ± 3.7%, N=3, n=12; *sema3fb^ca305^* 35.4 ± 3.0%, N=3, n=20; Mann Whitney U, p=0.45) and fractional shortening (WT 17.6 ± 2.0%, N=3, n=12; *sema3fb^ca305^*, 13.8 ± 1.6%, N=3, n=20; Mann Whitney U, p=0.17) (Figure 3H,I) were unchanged. Overall, the heart of the *sema3fb* mutant embryo was impaired significantly in function.

**Figure 3.**
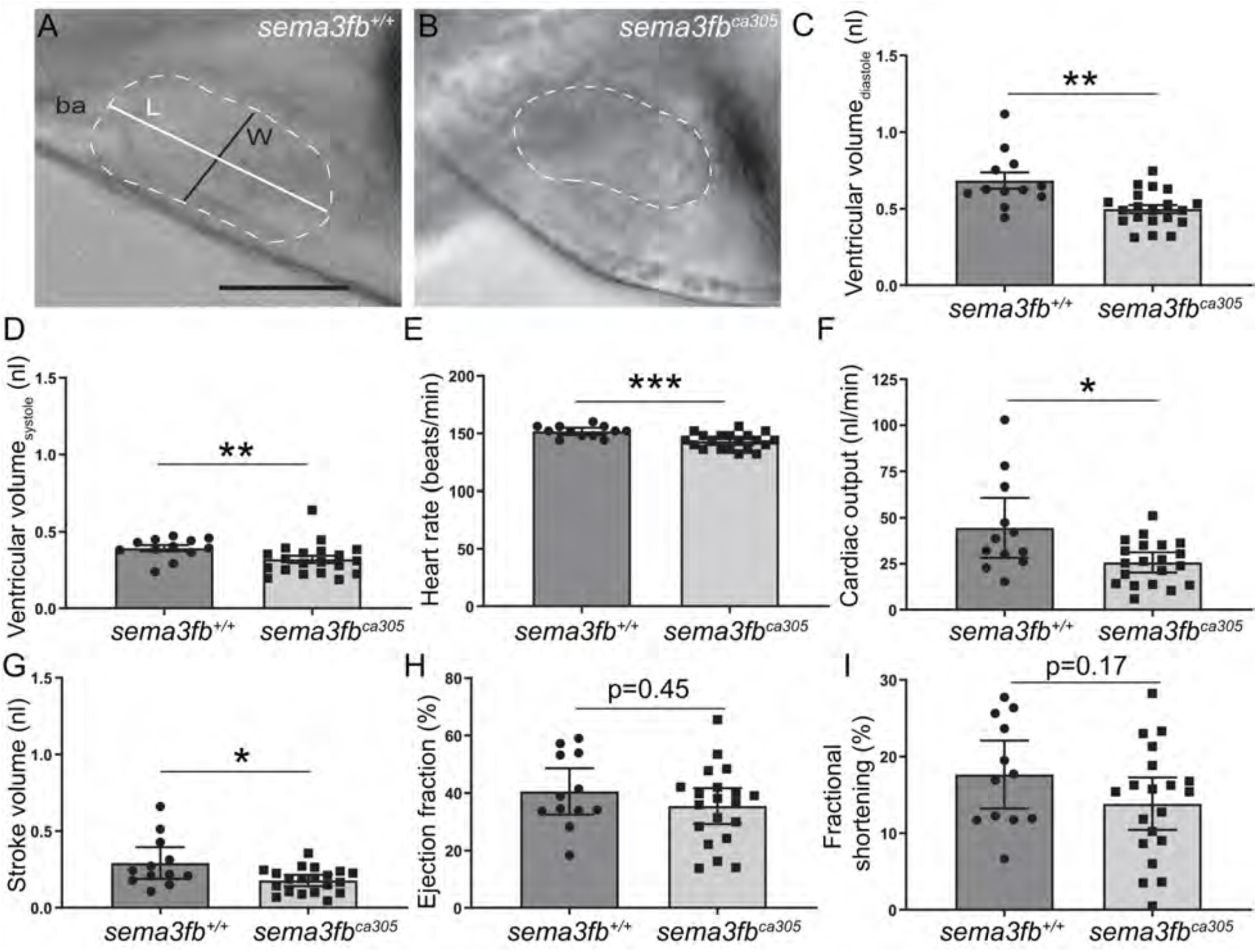
*sema3fb* mutant embryos have reduced cardiac function. **A-B)** Still images from videos of beating WT and *sema3fb^ca305^* mutant hearts at 72 hpf. Ventricle measurements were made as indicated, in both the long axis (white line, L: length) and short axis (black line, W: width) to determine ventricle volumes. ba: bulbus arteriosus. **C-D)** Quantitation of the average ventricle volume at end diastole (C) and end systole (D) reveals a significantly decreased capacity in the *sema3fb* mutant (**p=0.0027 and **p=0.0063, respectively) as compared to WT. **E-I)** Graphs showing average heart rate (***p=0.0008) (E), cardiac output (*p=0.026) (F), stroke volume (*p=0.033) (G), ejection fraction (p=0.45) (H), and fractional shortening (p=0.17) (I). Error bars are SEM over 3 independent experiments (n=12 *sema3fb*^+/+^, n=20 *sema3fb^ca305^*). Statistics represent the Mann-Whitney U test. Scale Bar: 100 µm.

### Early heart development occurs normally in the *sema3fb* mutants

The impaired function of the 72 hpf *sema3fb* mutant heart suggested a defect in heart development, which could include problems with cardiac progenitor cell specification and differentiation, morphogenesis of the heart tube, chambers and atrioventricular canal, formation of the endocardial heart valves, or the establishment of normal cardiac function^46^. To begin to address the underlying biology, we examined whether there were defects in early cardiac development, since *sema3fb* is expressed early in the bi-lateral heart fields (Suppl Figure 1B). Expression of *fgf8a*, a marker of the heart fields^47^, was not affected in the *sema3fb* mutants (Suppl Figure 2A-C), nor was its expression impacted elsewhere in the embryo, including in the brain, somites and tailbud region (Suppl Figure 2D-F), suggesting that early patterning was not disrupted by the *sema3fb* mutation. Next, we assessed the heart at 24 hpf by whole mount ISH for the cardiomyocyte marker *tnnt2a*, and for *myosin heavy chain 7* (*myh7*)^48^ and *myosin heavy chain 6* (*myh6*)^49^ to label ventricular and atrial cardiomyocytes, respectively (Suppl Figure 3). *sema3fb^ca305^* and *sema3fb^ca306^* hearts were generally in a cone/condensed state at 24 hpf (Suppl Figure 3B,C,F,G) and had not yet elongated into the linear tube of WT hearts^50, 51^ (Suppl Figure 3A,E,D,H,L). Nonetheless, two hours later tube formation had occurred in *sema3fb* mutants; 80% of *sema3fb^ca305^* (N=1, n=8/10) and 90% of *sema3fb^ca306^* (N=1, n=9/10) mutants exhibited elongated heart tubes at 26 hpf (Suppl Figure 3N,O), comparable to the numbers observed in WT siblings (N=1, n=9/10) at 23 hpf (Suppl Figure 3M,R). Notably, the cranial ganglia that emerge immediately adjacent to the heart developed normally in aged-matched embryos (Suppl Figure 3P,Q). Overall these data argue that heart morphogenesis is delayed only slightly (by 2-3 hours) by the loss of Sema3fb, with cardiomyocyte marker expression present at 24 hpf.

### Heart valve development disrupted slightly with loss of Sema3fb

Edema can result from a failure of valves to form between the chambers that results in retrograde flow of blood and fluid build-up^52^. Valves were present, however, when assessed in H&E-labeled 72 hpf *sema3fb^ca305^* heart sections (Figure 4A-B). To confirm these data, we bred the *sema3fb^ca305^* allele onto the *Tg(kdrl:mCherry)* line, in which the endocardial cells that make up the valves^52^ are mCherry positive at 72 hpf^25^. We visualized the heart by immunolabeling with the cardiac muscle marker MF20^53^. In both the WT and *sema3fb^ca305^* hearts, valves were present at the border between the atrium and ventricle (Figure 4C,D), and were of similar lengths (N=2; WT 61.6 µm ± 2.5 µm, n=10 embryos; *sema3fb^ca305^* 60.1 µm ± 3.8 µm, n=12 embryos; Mann Whitney U, p=0.72) (Figure 4J). The intensity of the mCherry label in the *sema3fb^ca305^* valves, however, was reduced as compared to WT (Figure 4C’,D’). Further, the distribution of the valves differed between the genotypes, with the valves in WT hearts roughly symmetrically distributed on either side of the constriction between the atrial and ventricular chambers, but preferentially located on the ventricular side of this border in mutants (Figure 4K).

**Figure 4.**
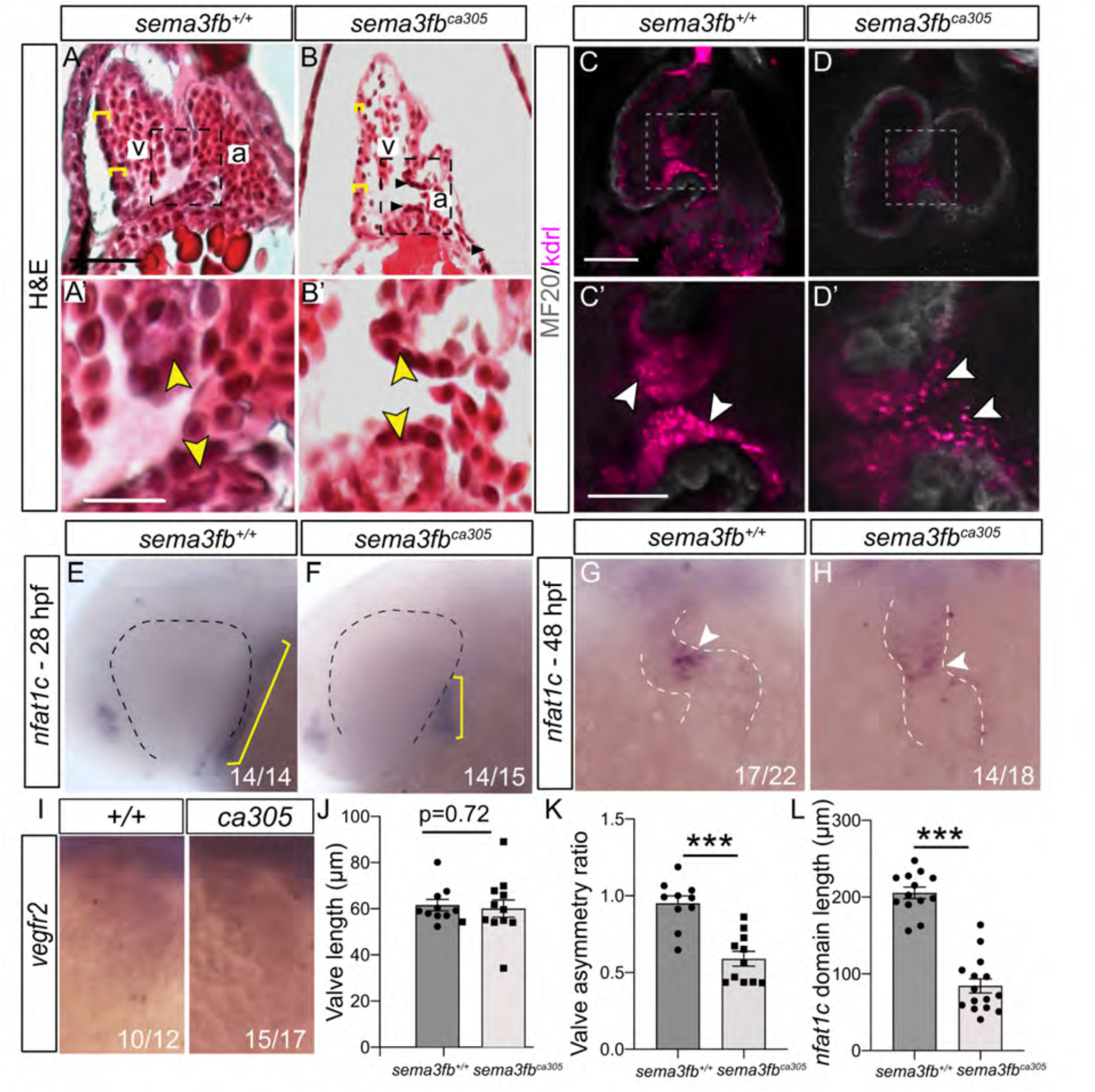
Disrupted heart valve development with the loss of Sema3fb. **A,B)** Histological sections stained by hematoxylin and eosin showing the ventricle walls (yellow brackets) and valves (arrowheads within inset) of WT (A, A’) and *sema3fb^ca305^* embryos (B, B’) at 72 hpf. **C-D)** Physical presence of the AV valve was confirmed in confocal projections of 72 hpf hearts immunolabeled by the myocardial marker MF20 (grey) on an endocardial-labeled transgenic line *Tg(kdrl:mCherry)* (arrowheads C’,D’) of WT (C, C’) and *sema3fb^ca305^* embryos (D, D’). **E-F)** Labeling in WT (E) and *sema3fb^ca305^* (F) embryos at 28 hpf reveals a smaller endocardial domain, as marked by *nfat1c* ISH label (yellow bars) in the mutant. **G-H)** By 48 hpf, the endocardial marker *nfat1c* is concentrated at the AVC (arrowheads) in WT (G) and mutant (H) fish. **I)** The endocardial marker *vegfr2* is expressed in the atrium and ventricle of both 48 hpf WT and *sema3fb^ca305^* fish. **J-L)** Quantitation in WT and mutant fish of the: (1) length of the AV valve, as marked by mCherry in the *Tg(kdrl:mCherry)* background (J; N=2, n=10 *sema3fb*^+/+^, n=12 *sema3fb^ca305^*), (2) the ratio of the length of the valve present in the atrium to that measured in the ventricle (K; N=2, n=10 *sema3fb*^+/+^, n=11 *sema3fb^ca305^*), and (3) the length of the *nfat1c* domain (marked by yellow bars in E and F) in 28 hpf WT and *sema3fb^ca305^* embryos (L; N=2, n=13 *sema3fb*^+/+^, n=15 *sema3fb^ca305^*). Error bars are SEM. Statistics represent the Mann-Whitney U test (***; p=0.0003 in K; p<0.001 in L).

To assess whether early development of the endocardium, from which valves form, occurred normally in the absence of Sema3fb, we examined the expression of the early endocardial marker *nfatc1*^54, 55^. While *nfatc1* ISH signal was present at comparable levels in WT and mutant hearts at 28 hpf, the domain in *sema3fb^ca305^* embryos (84.3 µm ± 9 µm, N=2, n=15 embryos) was significantly shorter than in WT (205.7 µm ± 7 µm, N=2, n=13; Mann Whitney U test, p<0.001) (Figure 4E,F,L). Yet, *nfatc1* mRNA was localized preferentially to the atrioventricular constriction at 48 hpf in both genotypes (N=2, WT, n=21/22 embryos; *sema3fb^ca305^*, n=16/18 embryos) (arrowhead, Figure 4G-H), and expression of the *vegfr2* endothelial cell marker was similar in the two genotypes (Figure 4I). These data suggest a delay in endocardial development, similar to what is seen with heart tube elongation (Suppl Fig. 3), but no gross defect in the initial formation of this tissue in *sema3fb^ca305^* embryos.

### Heart chamber size is decreased in *sema3fb* mutants

Heart chamber morphogenesis is complete by 48 hpf^50^, and thus a discernable defect in the morphology of the heart at this time-point would be expected to manifest as a functional deficit by 72 hpf. To determine the state of chamber development at 48 hpf we used the *tnnt2a* marker, which labels the entire heart. Looped hearts with a morphologically evident ventricle and atrium were present across all genotypes (Figure 5A-C). Measurements of the antero-posterior length of the *tnnt2a*-labelled heart (Figure 5D), however, revealed that the hearts of both *sema3fb^ca305^* (122.8 ± 5.0 µm, N=3, n=16; Kruskal-Wallis, Dunn’s multiple comparisons, p<0.0001) and *sema3fb^ca306^* (135.4 ± 4.5 µm, N=3, n=16; Kruskal-Wallis, p=0.0016) mutant embryos were significantly shorter than WT (163.8 ± 4.0 µm, N=3, n=18).

**Figure 5.**
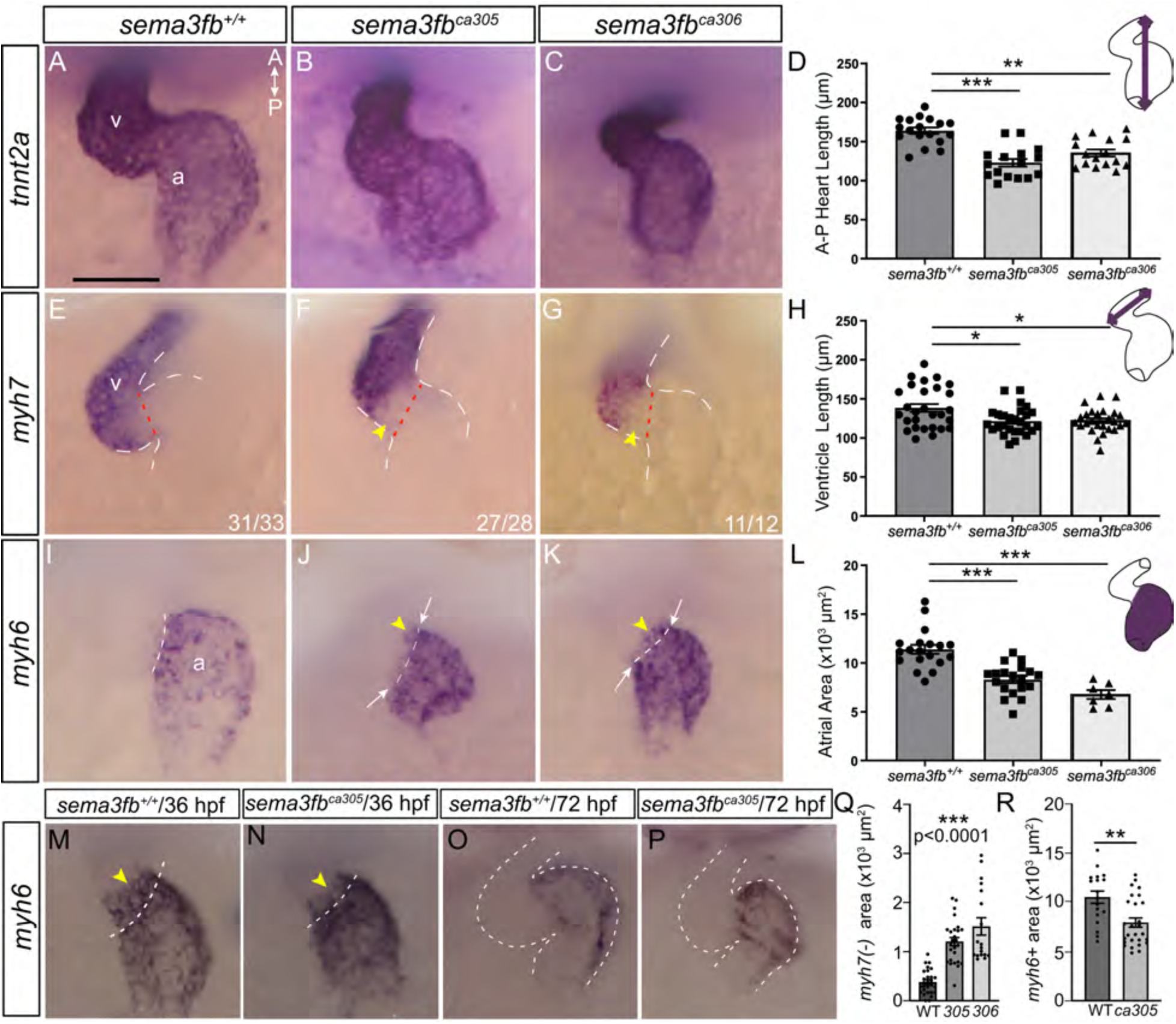
Ventricle and atrium are smaller in *sema3fb* mutant embryos. Ventral views of 48 hpf hearts labelled by whole mount ISH. Dotted lines mark the presumptive border between the atria and ventricle. **A-C)** The cardiomyocyte marker *tnnt2a* reveals total heart morphology. The WT heart loops normally and exhibits large chambers (N=2, n=16/18) (A), while both *sema3fb* mutant alleles present with normally looped, but smaller hearts (N=2, 14/16; N=2, 15/16) (B-C). **D)** Quantitation of the anterior to posterior length of the heart (purple line, schematic) reveals significantly smaller *sema3fb^ca305^* (N=3, n=16, ***p<0.0001) and *sema3fb^ca306^* (N=3, n=16, **p=0.0016) hearts as compared to WT (N=3, n=18). **E-G)** Both *sema3fb* mutant alleles (F,G) present with smaller *myh7* positive ventricles (N=2, 17/20 *sema3fb^ca305^*; N=2, 15/18 *sema3fb^ca306^*) than WT (N=2, n=19/20) (E). Arrows indicate ectopic *myh7* label in the atria of *sema3fb* mutants. **H)** Quantitation (purple line, schematic) reveals significantly shorter ventricles in *sema3fb^ca305^* (N=3, n=27, *p=0.009) and *sema3fb^ca306^* (N=3, n=26, *p=0.013) as compared to WT (N=3, n=28) embryos. **I-K)** Both *sema3fb* mutant alleles present with observably smaller *myh6* positive atria (N=2, 18/21 *sema3fb^ca305^*; N=2, 15/17 *sema3fb^ca306^*) (J-K) than in WT (N=2, n=19/19) (I). Asterisks indicate ectopic myh6 label in the ventricles of *sema3fb* mutants. **L)** Quantitation of surface area (purple fill) reveals significantly smaller atria in *sema3fb^ca305^* (N=3, n=19, ***p=0.0001) and *sema3fb^ca306^* (N=2, n=7, ***p<0.0001) hearts as compared to WT (N=3, n=19). (Kruskal Wallis One Way ANOVA, Dunn’s multiple comparisons test) **M-P)** *myh6* expression at 36 hpf (M,N), revealing expression anterior to the atrioventricular constriction (yellow arrowheads), and at 72 hpf (O,P). **Q)** Mean *myh7* negative area measured between the atrioventricular constriction and the posterior border of high *myh7* expression (N=3; WT n=28, *sema3fb^ca305^* n=26, *sema3fb^ca306^* n=18; p<0.0001, Kruskal Wallis One Way ANOVA, Dunn’s multiple comparisons test). **R)** Mean area of *myh6* ISH label at 72 hpf (N=2; WT n= *sema3fb^ca305^*; p=0.0021, Mann-Whitney U test). A: anterior; P: posterior. Scale bar: 50 µm.

To determine if one or both cardiac chambers were affected, we used whole mount ISH at 48 hpf for *myh7*^48^ and *myh6*^49^ to label ventricular and atrial cardiomyocytes, respectively. Both cardiac chambers were smaller in mutants as compared to WT. The long axis of the *sema3fb^ca305^* (121.5 ± 3.3 µm, N=3, n=27; Ordinary one-way ANOVA, Tukey’s multiple comparison test, p=0.009) and *sema3fb^ca306^* (122 ± 3.1 µm, N=3, n=26; Ordinary one-way ANOVA, p=0.013) ventricles was significantly shorter than in WT (138.4 ± 5.1 µm, N=3, n=28) (Figure 5E-H), as confirmed by measurements of the ventricle length during diastole (WT 203.9 ± 6.7 µm, n=12; *sema3fb^ca305^* 190.5 ± 2.3 µm, n=20; Mann Whitney U, p=0.048) and systole (WT 175.6 ± 4.2 µm, n=12; *sema3fb^ca305^* 163.3 ± 3.4 µm, n=20; Mann Whitney U, p=0.031) in live embryos viewed in brightfield (Suppl Figure 4). Additionally, the *myh6*+ atria were significantly smaller in area in the *sema3fb^ca305^* (8,290 ± 360 µm^3^, N=3, n=19; Kruskal-Wallis, p=0.0001) and *sema3fb^ca306^* (6,760 ± 450 µm^3^; N=2, n=7 Kruskal-Wallis, Dunn’s multiple comparisons, p<0.0001) hearts than in WT (11,390 ± 460 µm^3^; N=3, n=19) (Figure 5I-L). Ultimately, these data suggest that with the loss of Sema3fb both ventricle and atrium are specified, but the chambers are smaller in size than in their WT counterparts.

### The border between the ventricle and atrium appears disrupted with loss of Sema3fb

Interestingly, close inspection revealed that borders of the domains of expression of atrial and ventricular specific *myosins* at the atrioventricular junction were disrupted in *sema3fb* mutants. In the vast majority of WT hearts (N=4, 31/33 hearts), the *myh7*-expression border aligned with the constriction between the ventricle and atrial chambers (Figure 5F). In contrast, in both the *sema3fb^ca305^* (N=4, n=27/28 hearts; Figure 5F) and *sema3fb^ca306^* (N=2, n=11/12; Figure 5G) heart, expression of *myh7* was low or absent from the atrioventricular band (yellow arrowheads, Figure 5F,G), such that the posterior border of the high *myh7* expression domain sat anterior to the band. We quantitated this phenotype by measuring the *myh7* negative area between the atrioventricular constriction and the posterior domain of solid *myh7* expression (Figure 5Q), and found a significantly larger non/low-expressing domain in both mutants than in WT (WT 383 ± 46 µm^3^, n=28 embryos; *sema3fb^ca305^* 1205 ± 86 µm^3^ n=26; *sema3fb^ca306^* 1516 ± 177 µm^3^, n=18; p<0.0001 Kruskal-Wallis test, Dunn’s multiple comparisons). Additionally, while the border of the *myh6* atrial marker expression domain was generally smooth in WT hearts (N=3, n=22/26 hearts; Figure 5I), it was often disorganized in both the *sema3fb^ca305^* (N=3, n=23/31) and *sema3fb^ca306^* (N=1, n=4/7) hearts (Figure 5J,K). Furthermore, in most WT hearts the anterior border of *myh6* expression aligned with the morphological constriction between the atrium and ventricle (n=20/26), but often extended a small extent past this border and into the ventricle in *sema3fb^ca305^* (N=4, n=23/31) and *sema3fb^ca306^* (N=1, n=6/7) hearts (Figure 5I-K). Of note, in younger 36 hpf hearts this antero-wards extension of the *myh6* domain past the constriction was present in both WT and *sema3fb^ca305^* genotypes (Figure 5M,N), suggesting that there was a failure to resolve to the mature border in mutants. The defect in chamber size was still present in *sema3fb* mutants at 72 hpf, as evident by *myh6* expression in the atria (Figure 5O,P), and confirmed by quantitation of the area of the atrial *myh6*+ expression domain in WT vs. mutants (N=2; WT, 10,518 ± 618 µm^3^, n=18 embryos; *sema3fb^ca305^* 7929 ± 472 µm^3^, n=27; p=0.0021 Mann Whitney U test) (Figure 5R). These data indicate that with Sema3fb loss there is disruption of the expression of mRNA for atrial and ventricular myosins at the border between the two chambers. We also asked if expression of a non-myosin ventricular chamber specific gene was disrupted at the atrioventricular junction. Normally, *tnnt2a* mRNA is expressed at higher levels in the ventricle than atrium, as evidenced in WT hearts by a sharp drop in mRNA levels in the atrium relative to the ventricle (Figure 6A). This drop was less dramatic in both *sema3fb* mutant alleles, and an obvious border between expression in the two chambers was lacking.

**Figure 6.**
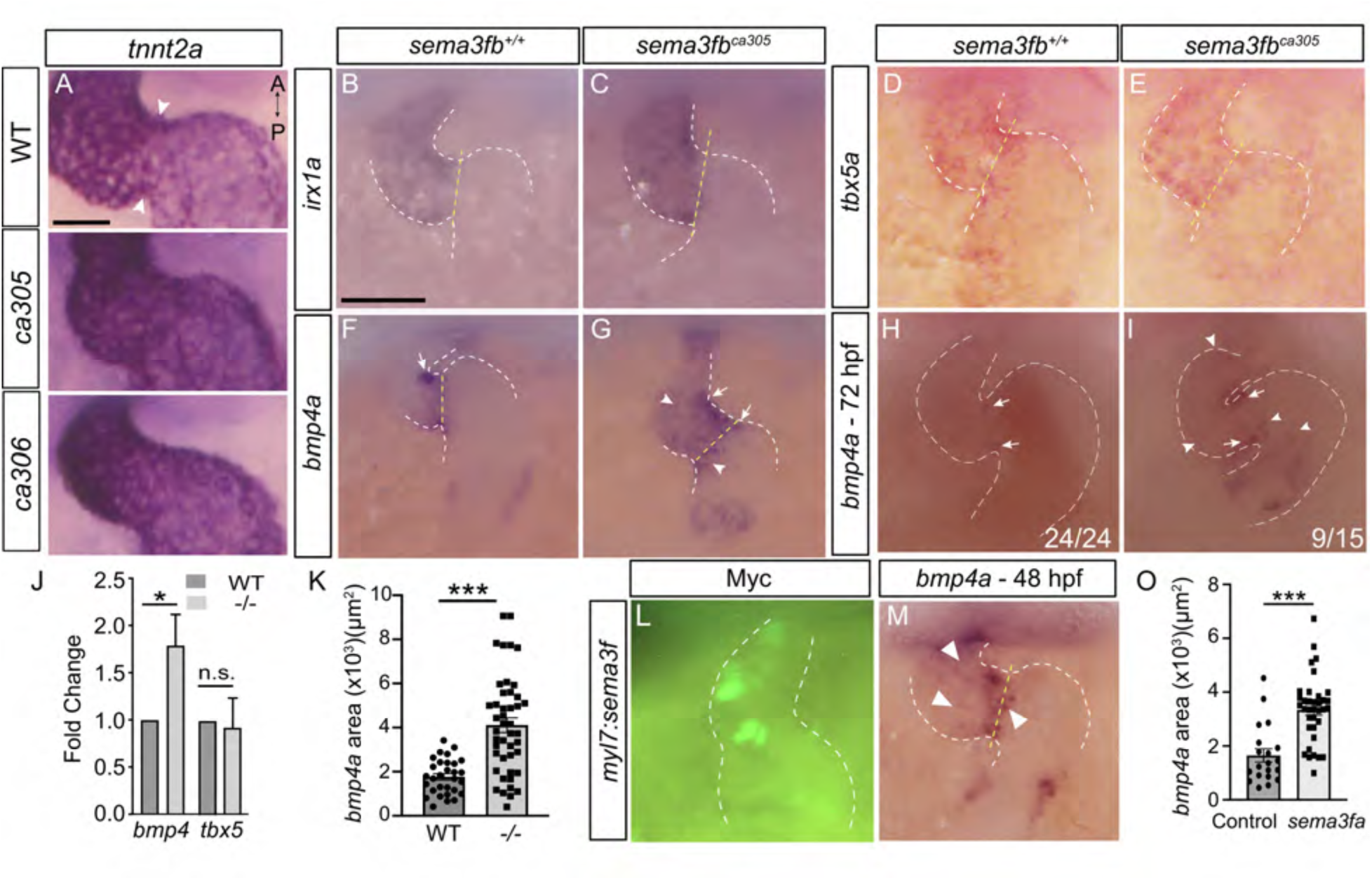
Chamber identity disrupted at the AVC with loss and overexpression of Sema3f. Ventral images of 48 hpf (A-G) whole mount *sema3fb*^+/+^ and *sema3fb^ca305^* embryos processed for ISH. **A)** The cardiomyocyte *tnnt2a* marker does not show the sharp expression change at the border between the atrium and ventricle in the *sema3fb^ca305^* mutant heart that is present in WT (white arrowheads). **B-E)** Expression of the ventricle markers *irx1a* (B-C; N=3; WT n=15/15; *sema3fb^ca305^* n=16/16) and *tbx5a* (D-E; N=1; WT n=9/9; *sema3fb^ca305^* n=7/7) are unchanged in *sema3fb^ca305^* as compared to WT hearts. **F-I)** Expression of *bmp4a* is restricted to the AV valves (white arrow) in WT hearts at 48 hpf (F, N=4, 70%, n=21/30) and 72 hpf (H, N=1, n=8/9), but expands (white arrowheads) into the ventricle and/or atria of mutants (G, 48 hpf, 80%, n=36/45 with expansion; I, 72 hpf, n=10/11). **J)** RT-qPCR reveals significant upregulation of *bmp4a* (*p=0.036, N=5) and no change in *tbx5a* (p=0.7, N=3) mRNA levels in *sema3fa^ca305^* mutant vs. WT embryos. Error bars represent standard deviation (SD), and the statistics represent the Mann-Whitney U test. **K)** Graph of the mean area of the *bmp4a* domain measured from images of whole mount *bmp4a* ISH (N=4, WT n=30, mutant n=45; Mann-Whitney U test, p<0.0001). **L-O)** Ventral images of 48 hpf whole mount embryos injected at the one cell stage with *myl7:sema3f* plasmid to drive Sema3f expression in heart cardiomyocytes, and processed by immunolabeling for myc to visualize the myc-tagged Sema3f (L), and by ISH for *bmp4a* (M). With Sema3f overexpression *bmp4a* expands into the ventricle and/or atria (white arrowheads, J; N=2, n=24/37 expanded) as compared to control (*bmp4a* n=17/20 normal). **O)** Graph of the mean area of the *bmp4a* domain measured from whole mount ISH mages (N=2, control n=20, Sema3f overexpression n=36; Mann-Whitney U test, p<0.001). Scale bar in A (panels A-C) and B (panels B-I, L-M) is 50 µm.

Possibly, the disrupted chamber-specific myosin expression in mutants was a feature of altered chamber-specific cardiomyocyte differentiation. Thus, we looked at the expression of the ventricle-specific homeobox-containing transcription factor, *iroquois1a* (*irx1a*)^56^ at 48 hpf. Interestingly, despite the reduction of ventricular marker *myh7* mRNA at the atrioventricular junction, *irx1a* mRNA expression in mutant and WT hearts was similar, with expression extending through the atrioventricular canal in both genotypes (Figure 6B,C; N=2; WT, n=15/15 hearts; *sema3fb^ca305^* n=16/16). In agreement, the high-to-low ventricle-to-atrium gradient of *T-box transcription factor tbx5a*^57^ mRNA was unchanged between WT (N=1, n=9) and *sema3fb^ca305^* (N=1, n=7) embryos (Figure 6D,E), as verified by RT-qPCR (-/- 0.93 ± 0.3% (standard deviation) fold change vs. WT, N=3; Mann Whitney U, p=0.7; Figure 6I). These data indicate that the ventricular differentiation program is not affected globally by Sema3fb loss.

Because of the altered expression of *myh7* at the AVC, we next asked whether *bone morphogenetic protein 4a* (*bmp4a*) expression at the AVC was regulated appropriately. *bmp4a* is expressed initially throughout the antero-posterior length of the myocardium and then becomes largely restricted to the AVC by 37 hpf^58^. By 48 hpf, we found the strongest *bmp4a* mRNA signal was in the AVC region in almost all WT (N=4; 96.7%, n=29/30), and *sema3fb^ca305^* (N=4; 91%, n=41/45) hearts (Figure 6F,G). In the majority of WT hearts, *bmp4a* was restricted to the AVC alone (N=4; 70%, n=21/30), with a smaller number of hearts showing additional ISH signal throughout the ventricle (20% n=6/30) or in the atrium (13.3%, n=4/30). In contrast, in mutant hearts *bmp4a* mRNA often continued to be expressed throughout the ventricle (71.1%, n=32/45) and/or in the atrium close to the AVC (53.3% n=24/45). RT-qPCR verified that at 48 hpf *bmp4a* levels were upregulated in the *sema3fb^ca305^* hearts (-/- 1.79 ± 0.33% (standard deviation) fold change vs. WT, N=5; Mann Whitney U, p=0.036) (Figure 6J). Furthermore, quantitation revealed a significantly larger *bmp4a* expression domain in *sema3fb* mutant hearts than WT (Figure 6K; N=4; WT 1770±145 µm^3^, n=30; *sema3fb^ca305^* 4109±332 µm^3^ n=45; p<0.0001, Mann-Whitney U test). By 72 hpf, mutant hearts still exhibited a handful of *bmp4a*- expressing ectopic cells within the ventricle (60% n=9/15) or atrium (73% n=11/15), though *bmp4a* expression was concentrated mainly at the AVC, as was the case in all WT hearts (n=24/24) (Figure 6H,I).

Sema3f may be required either to inhibit expression of *bmp4a* throughout the ventricle or to mark the border between atrium and ventricle and identify the domain to which *bmp4a* expression should resolve. We used Sema3f overexpression to address these two possibilities. If the former was true, we expected to see a knockdown of *bmp4a* levels in hearts overexpressing Sema3f. In the latter scenario, misexpression of Sema3f would disrupt the border and a broadened *bmp4a* AVC expression domain might result. To overexpress Sema3f in developing cardiomyocytes we injected a Tol2 *myl7:sema3f-myc* construct into embryos at the one-cell stage along with *transposase* mRNA, with injection of the plasmid alone serving as control. We tried but failed to clone *sema3fb*, however, a full-length cDNA clone of zebrafish *sema3fa* was available. Sema3fa and Sema3fb paralogs share 83% amino acid identity, and *sema3fa* or *sema3fb* mRNA both result in delayed neutrophil recruitment in a zebrafish model of inflammation^59^. Thus, we expected Sema3fa to behave like Sema3fb. At 48 hpf, the *sema3f*- expressing fish were processed by whole mount ISH for *bmp4a* mRNA. Notably, Sema3f was expressed in a mosaic fashion in both the atrium and ventricle, as visualized by immunolabeling for the myc-tagged Sema3f (Figure 6L). Sema3f overexpression produced a similar *bmp4a* phenotype to that seen with loss of Sema3fb. The control and Sema3f-expressing hearts both exhibited enhanced expression of *bmp4a* at the AVC by 48 hpf (Control, 100% n=20/20; Sema3f overexpression, 97% n=35/36), however, *bmp4a* ISH label expanded beyond the AVC with Sema3f overexpression (arrowheads, Figure 6M). *bmp4a* mRNA was present only within the AVC in most control hearts (75%, n=15/20), but this was true in a small number of *sema3f*- overexpressing hearts (25%, n=9/36), where the *bmp4a* domain spread beyond the AVC into the atrium and/or ventricle. In support, the area of the *bmp4a* mRNA domain was significantly smaller in the Sema3f-overexpressing hearts than control (Figure 6O). These data argue that Sema3fb does not directly regulate *bmp4a* expression, but that *bmp4a* expression is altered as a downstream consequence of the loss of Sema3fb signaling. The fact that both loss and gain of Sema3f function produced a similar phenotype argues that normally Sema3fb signaling needs to be strictly regulated, and that either too little or too much signaling disrupts normal development of the border between the ventricular and atrial chambers.

To confirm at the protein level that there was disruption of the border between the two heart chambers in *sema3fb* mutants, we used two antibodies that label proteins that are expressed at higher levels by ventricular than atrial myocardium at 48 hpf: MF20 labels cardiac muscle and is commonly used as a ventricular marker in zebrafish^57, 58^, and an antibody against the cell-surface adhesion molecule DM-GRASP that labels the surface of cardiomyocytes^60^. Whole mount immunostaining, followed by confocal microscopy, revealed clearly separate atria and ventricles in WT hearts at 48 hpf (N=1, n=5/5 MF20; N=3, n=11/12 DM-GRASP, Figure 7A,C). In contrast, the distinct border between the chambers was disrupted in at least 83% of mutants (N=1, n=8/9 MF20; N=3, n=10/12 DM-GRASP, Figure 7B,D). This loss of a sharp ventricle to atria border was evident with the quantitation of the average intensity of MF20 immunolabel over the antero-posterior axis of the heart (WT n=5, *sema3fb^ca305^* n=9; Figure 7E).

**Figure 7.**
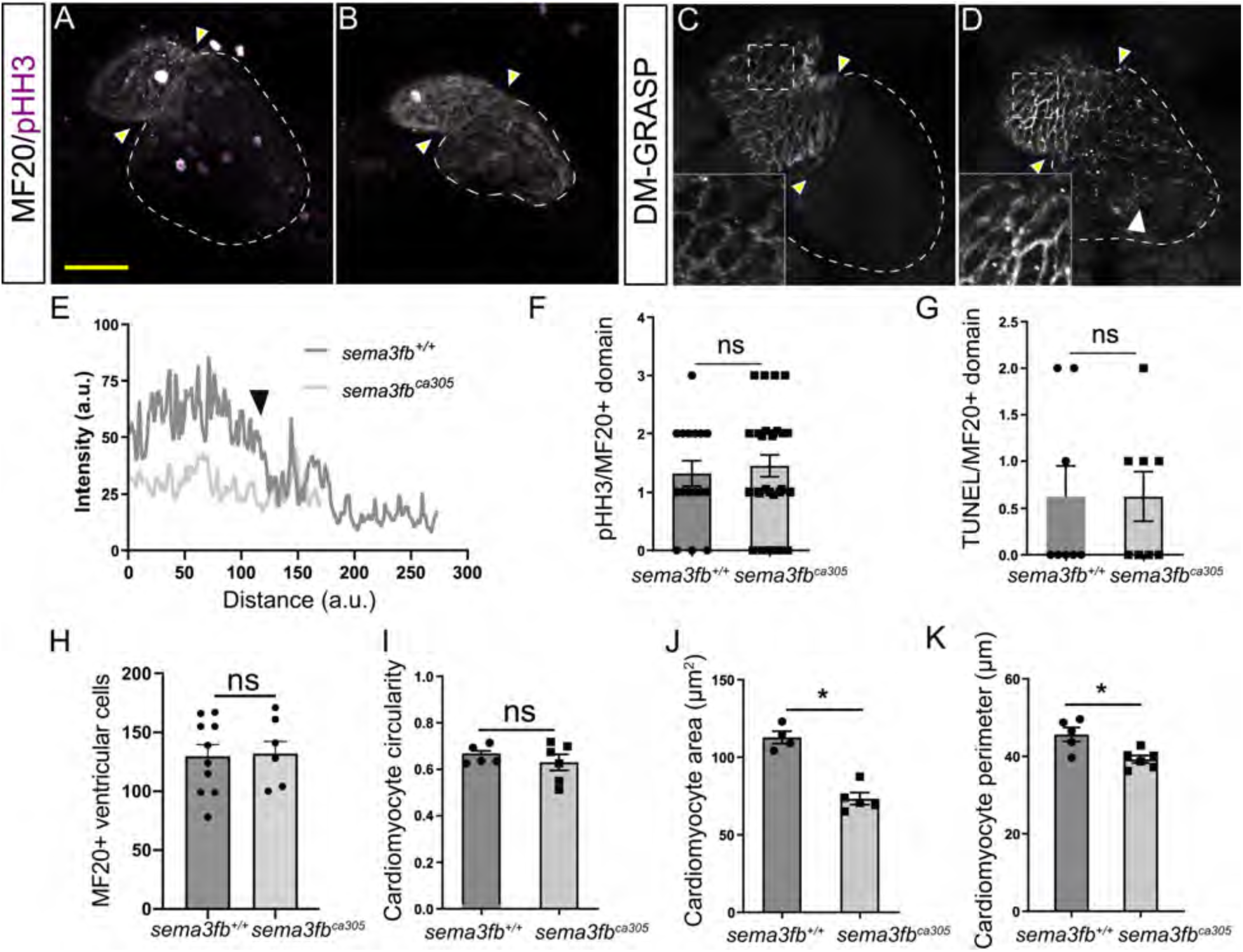
Loss of Sema3fb disrupts the normal pattern of ventricular cardiomyocyte protein expression. **A-D)** Confocal images of 48 hpf whole mount hearts processed by immunohistochemistry for pHH3 and MF20 (A,B), and DM-GRASP (C,D). MF20 and DM-GRASP are more highly expressed in the WT ventricle, with protein expression exhibiting a fairly sharp border at the atrioventricular constriction (yellow arrowheads) (A, N=1, n=5/5, and C, N=3, n=11/12). Expression of both proteins expands into the atrium of the mutant hearts (B, N=1, n=8/9 and D (white arrowhead), N=3, n=10/12). **E)** Average RGB immunolabel intensity for MF-20 across the antero-posterior axis of the heart reveals that WT (n=5) hearts exhibit high expression in the ventricle that sharply tapers at the border with the atrium (∼150 arbitrary units (a.u.); arrowhead), while the intensity profile in *sema3fb^ca305^* (n=9) mutants remains stable through both chambers. Note that hearts are smaller in the *sema3fb* mutants. **F)** Mean number of actively dividing MF20+ cardiomyocytes as revealed by pHH3 immunostaining (WT n=16 hearts, mutant n=27, p=0.71). **G)** Quantitation of the number of TUNEL positive MF20-expressing cardiomyocytes at 48 hpf (N=2, n=8, both genotypes, p=0.99). **H)** Mean number of MF20+ ventricular cardiomyocytes in DAPI-labelled hearts (n=10 WT and n=7 mutants; p=0.72). **I-K)** Mean ventricular cardiomyocyte area (J) and perimeter (K) are significantly smaller in *sema3fb^ca305^* as compared to WT (p=0.016 and p=0.03, respectively), while circularity (I) is not impacted (p=0.66). Measurements were made using N=3, n=4-5 WT and N=3, n=6 mutant embryos, with 9-12 DM-GRASP labelled cells measured per embryo. Error bars represent SEM. Statistics represent the Mann-Whitney U test. A: anterior; P: posterior. Scale bar: 50 µm.

### A decrease in cardiomyocyte size explains the small ventricles observed in *sema3fb* mutants

The small size of the ventricles and atria of *sema3fb* mutants at 48 hpf could be due either to fewer cardiomyocytes or a reduction in cardiomyocyte size. While our data indicate that cardiomyocyte progenitor cells are specified and differentiate in the absence of Sema3fb, subsequently less proliferation and/or increased apoptosis could reduce their numbers. Similar numbers of mitotically active cardiomyocytes, however, were labelled with an antibody against phosphorylated histone H3 (pHH3) in 48 hpf WT and *sema3fb^ca305^* hearts (WT 1.31 ± 0.2 cells, N=3, n=16; *sema3fb^ca305^* 1.44 ± 0.2 cells, N=3, n=27; Mann Whitney U, p=0.71) (Figure 7A,B,F). Furthermore, no significant changes were observed at 48 hpf in the numbers of TUNEL positive apoptotic cardiomyocytes in WT and *sema3fb^ca305^* embryos (WT 0.62 ± 0.3 cells, N=2, n=8; *sema3fb^ca305^* 0.62 ± 0.3 cells, N=2, n=8; Mann Whitney U, p>0.99) (Figure 7G). In support of these data, similar numbers of MF20+ cells in the ventricle were observed in WT and mutant hearts at 48 hpf (Figure 7H).

Cardiomyocytes normally begin to enlarge as the zebrafish cardiac chambers emerge (36 hpf), and between 40–45 hpf more than half of ventricular cardiomyocytes increase in size^60, 61^. Thus, we asked whether cardiomyocyte size was impacted in the mutants, and could explain the decreased heart size at 48 hpf. We focused on ventricular cardiomyocytes, because in WT hearts DM-GRASP immunostaining labeled ventricular cardiomyocyte membranes, but not to any great extent atrial cells (Figure 7C). Interestingly, the size of ventricular cardiomyocytes was strikingly different between the WT and *sema3fb^ca305^* embryos (insets, Figure 7C,D). To quantitate this change, we took confocal projections of DM-GRASP-labeled hearts and measured area, perimeter and circularity of the cardiomyocytes within the central region of the ventricle. While circularity was comparable across WT and mutant cardiomyocytes (WT 0.66 ± 0.02, n=5 embryos (9-12 cells measured/embryo); *sema3fb^ca305^* 0.63 ± 0.03, n=6 embryos; Mann Whitney U, p=0.66, Figure 7I), both the area (WT 112.8 ± 4.0 µm^3^, n=4 embryos; *sema3fb^ca305^* 73.3 ± 3.8 µm^3^, n=5 embryos; Mann Whitney U, p=0.016, Figure 7J) and perimeter (WT 45.6 ± 1.8 µm, n=5 embryos; *sema3fb^ca305^* 39.2 ± 1.0 µm, n=6 embryos; Mann Whitney U, p=0.03, Figure 7K) of ventricular cardiomyocytes were reduced significantly in the mutant embryos. These data support the idea that chamber size is decreased in mutants in part because of a reduction in cardiomyocyte size.

A significant portion of the cardiac myocyte population in zebrafish arises from the integration of cardiomyocytes from the secondary heart field^62^ and cardiac neural crest (CNC)^63^. To determine if cardiomyocytes originating from either of these two sources failed to enter the heart as a result of the loss of a known chemotropic molecule expressed therein, we assayed markers of the secondary heart field (*latent tgfβ binding protein 3a* (*ltbp3a*))^64, 65^ and CNC (*cysteine-rich intestinal protein 2* (*crip2*)) by whole mount ISH at 48 hpf. *crip2* was expressed at similar levels in WT and mutant ventricles (Suppl Figure 5A,B), and *ltbp3a*-expressing cardiomyocytes were present in both the WT and mutant hearts (Suppl Figure 5C,D). Together, our data argue that changes in proliferation, apoptosis, and the integration of cardiomyocytes from sources external to the heart, do not contribute substantially to the small stature of the *sema3fb* mutant heart.

Because cardiomyocyte size is linked intimately to the hemodynamic forces of flow within the heart^60, 61^, we asked whether *sema3fb* mutant hearts were still smaller than their WT counterparts when cardiac flow was removed from the equation by growing hearts independent of the circulatory system. Hearts of WT and mutant embryos were explanted at 24 hpf and cultured for 24 hours. Explants were fixed and assessed by ISH for the expression of atrial *myh6* (Suppl Figure 6A,B). *myh6*+ mutant atria were significantly smaller (WT 2950 ± 210 µm^3^, N=2, n=8 explants; *sema3fb^ca305^* 2320 ± 130 µm^3^, N=2, n=13 explants; Mann Whitney U, p=0.025) in size as compared to WT (Suppl Figure 6C), arguing that chamber defects present in the Sema3fb mutants independently of hemodynamic forces.

### Nrp2b and Plxna3 receptors may mediate the Sema3fb signal

Sema3s signal canonically through Nrp and Plxn receptors^13^. We next asked which receptors Sema3fb might act through to influence cardiac chamber development by performing whole mount ISH for *plxn* and *nrp* genes in zebrafish. Expression of zebrafish *nrp* genes has been reported previously^67, 68^, and analysis of gross defects with Nrp knockdown, revealed vascular issues in *nrp1a*, *nrp1b* and *nrp2a* antisense MO-injected fish, and pericardial defects in the Nrp2b morphants^68^, pointing to a role for Nrp2b in heart development. *nrp2b* and *plxna3* mRNAs were present early alongside *sema3fb* mRNA in the bilateral heart field at 18 hpf (Suppl Figure 7A-C), with mRNAs confined to the presumptive ventricular myocardium by 24 hpf (Suppl Figure 7D; Figure 8B), as evident for *plxna3* at 48 hpf (Figure 8C). Of note, other *plxna* genes were not expressed at significant levels in heart tissue at 24 hpf (Suppl Figure 7H-L). In contrast, *sema3fb* mRNA was expressed throughout the entire myocardium (Figure 8A). To identify which heart cells express *plxna3* and *nrp2b* and can respond to Sema3fb we performed whole mount ISH on *Tg(flk:EGFP)* fish at 28-36 hpf. In these fish, endocardial cells in the heart are fluorescently labeled. For both *nrp2b* (Suppl Figure 7E-E’’) and *plxna3* (Figure 8D-D’’), ISH label was in cardiomyocytes adjacent to the EGFP-expressing endocardium. In summary, while *sema3fb* is expressed throughout the embryonic ventricle and atrial myocardium, the receptor expression data argue that Sema3fb signaling is likely spatially restricted to the ventricular myocardium.

**Figure 8.**
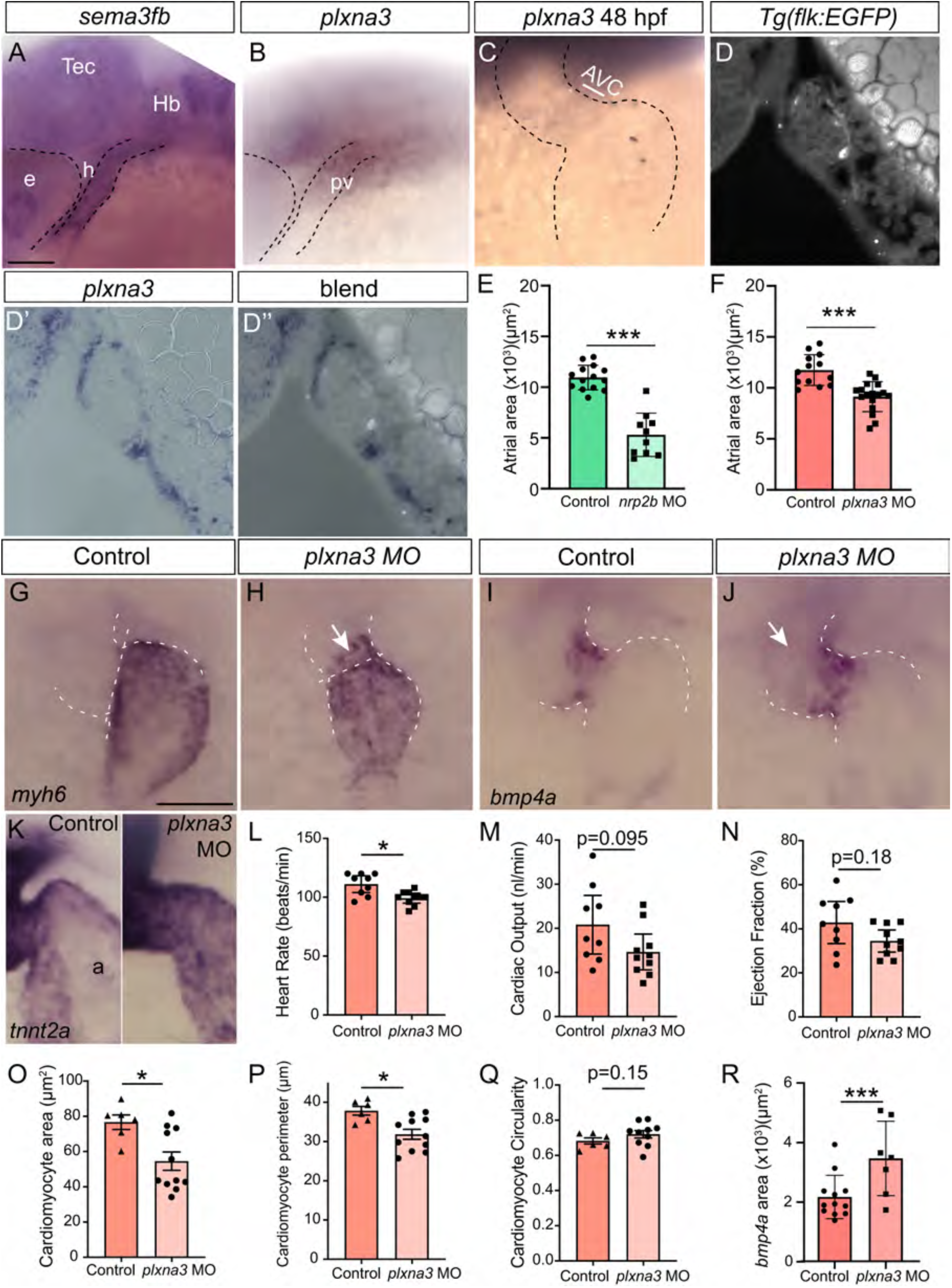
Ventricular expression of *plxna3* may restrict Sema3fb signaling to the ventricle. **A-C)** Lateral view of whole mount ISH for *sema3fa* (A) and *plxna3* (B) at 28 hpf, and a ventral view of *plxna3* mRNA in the ventricle but not atrium at 48 hpf (C). **D)** Sagittal section of *plxna3* (D’) ISH label at 36 hpf on a *Tg(flk:EGFP)* background to label the GFP+ endocardium (D). The blend of these two labels is shown in D’’. e, eye; h, heart; Hb, hindbrain; pv, presumptive ventricle; Tec, optic tectum. **E-F)** Mean atrial area is reduced significantly in *nrp2b* deficient embryos (p<0.0001, n=10 embryos; E) as compared to controls (n=14), and in *plxna3* knockdown embryos (N=2; p<0.0001, n=17; F) as compared to controls (n=13). **G-H)** Representative images of *myh6*-expressing atria of control (G) and *plxna3* (H) morpholino-injected embryos. White arrow delineates spillover of ISH label into the ventricle. **I-J)** Representative images of *bmp4a* expression in control (I) and *plxna3* (J) morpholino-injected embryos. White arrow and arrowhead delineate spillover of ISH label into the ventricle and atria, respectively. **K**) The cardiomyocyte label *tnnt2a* shows a sharp reduction in the atrium of a control but not a *plxna3* MO embryo. **L-N)** Graphs showing average heart rate (*p=0.011, L), cardiac output (p=0.095, M), and ejection fraction (p=0.18, N) (N=3, n=9 controls, n=10 plxna3 morphants). **O-Q)** Quantitation of cell size of DM-GRASP-labelled ventricular cardiomyocytes. A significant reduction is seen in cardiomyocyte area (O, *p=0.019) and perimeter (P, *p=0.015) in *plxna3*-deficient embryos as compared to controls, but not circularity (Q, p=0.15). Measurements were made using N=3, n=6 control and N=3, n=11 *plxna3* morphant embryos, with 8-10 cells per embryo. **R)** Average area of the *bmp4a* expression domain in control (n=12; p=0.028) and *plxna3* morphants (n=7). Error bars are SEM. Statistics represent the Mann-Whitney U test. Scale bar in A: 50 µm (A,B,D-D’’,K), 25 µm (C,G-J).

To determine whether loss of function of either receptor recapitulated the *sema3fb* mutant chamber phenotypes, we performed MO-mediated knockdown of Plxna3 and Nrp2b with previously characterized MOs^37^. Knockdown of either receptor resulted in obvious edema by 48 hpf, indicating that heart function was compromised. The hearts of *nrp2b*-deficient embryos were grossly malformed, with a severe reduction in atrial size (control 10,960 ± 320 µm^3^, n=14; *nrp2b* MO 5,314 ± 670 µm^3^, n=10; Mann Whitney, p=<0.0001, Suppl Figure 7F,G, Figure 8E), suggesting a key role for Nrp2b in early cardiac development. This early role meant, however, that we could not easily assess Nrp2b function in later aspects of heart development. Thus, we restricted our analysis to the *plxna3*-deficient embryos. Importantly, similar to what we saw in *sema3fb* mutants, *plxna3* morphants (Figure 8H) exhibited significantly smaller sized atria than control (Figure 8G) (N=2; control 11,749 ± 420 µm^3^, n=13; *plxna3* MO 9,142 ± 355 µm^3^, n=17; Mann Whitney U, p=<0.0001, Figure 8F). We next assessed heart function by live imaging of 72 hpf hearts. Similar to *sema3fb^ca305^* embryos, *plxna3*-deficient embryos showed a reduction in heart rate (99.2 ± 1.9 beats/min, N=3, n=10; Mann Whitney, p=0.011, Figure 8L) as compared to controls (111.1 ± 3.1 beats/min, N=3, n=9). Yet, cardiac output (control 20.8 ± 2.9 nL/min, N=3, n=9; *plxna3* MO 14.6 ± 1.8 nL/min, N=3, n=10; Mann Whitney, p=0.09; Figure 8M) and ejection fraction (control 42.9 ± 4.1%, N=3, n=9; *plxna3* MO 34.4 ± 2.2%, N=3, n=10; Mann Whitney, p=0.18; Figure 8N), were not significantly different between morphant and control hearts, despite the clear heart function defect revealed by the presence of edema in the *plxna3* morphants (data not shown). Likely the partial loss-of-function that would be generated with the *plxna3* MO produced cardiac defects that were less severe than those seen in the *sema3fb* mutant, but that over time resulted in edema.

Thus, it was important to determine whether in addition to the smaller heart, the key alterations observed in chamber development present in *sema3fb* mutants were also present to some degree in the *plxna3* morphants. Importantly, the area of individual DM-GRASP immunolabeled ventricular cardiomyocytes (control 76.7 ± 4.1 µm^3^, N=2, n=6 hearts; *plxna3* MO 54.6 ± 5.2 µm^3^, N=2, n=11; Mann Whitney U, p=0.019; Figure 8O), and their perimeter (control 37.8 ± 1.1 µm, N=2, n=6; *plxna3* MO 31.9 ± 1.3 µm, N=2, n=11; Mann Whitney U, p=0.014; Figure 8P), were significantly smaller in morphants than controls. Circularity of the morphant cardiomyocytes was comparable to controls (control 0.68 ± 0.02, N=2, n=6; *plxna3* MO 0.72 ± 0.02, N=2, n=10; Mann Whitney U, p=0.15; Figure 8Q). Notably, the distinct border of gene expression between the ventricle and atrium was also less evident in the *plxna3* morphants, as seen by the presence of strong *tnnt2a* expression in the atria (Figure 8K) and ectopic *bmp4a* expression in the ventricle in addition to the expected AVC labeling (Figure 8I,J,R). Together, these data argue that Plxna3 functions downstream of Sema3fb in the developing cardiac system. The *plxna3* morphant heart is impacted less severely than that of the *sema3fb* mutant, presumably because of residual Plxna3 signaling in the morphants.

## Discussion

Our data argue that embryonic cardiomyocytes secrete Sema3fb that spatially restricts chamber-specific gene expression at the atrioventricular border to ensure proper function of the embryonic heart. We find *sema3fb* mRNA is expressed by all cardiomyocytes over the period of chamber morphogenesis and differentiation, while expression of mRNAs for the candidate receptors, Nrp2b and Plxna3, is spatially restricted to ventricular cardiomyocytes. We propose that Sema3fb signals inform ventricular cardiomyocytes they border cells in the atrium, which lack the machinery to sense Sema3fb. The consequence is the proper refinement of chamber-specific gene expression at the AVC, a feature that is disrupted in the absence of Sema3fb. The size of both heart chambers is smaller, we propose as a consequence of these defects in gene expression. Our work argues for the first time that a heart-derived secreted Sema3 provides a spatially-restricted signal required for the normal refinement of the border between the ventricle and atrium, and that is required for proper chamber specific differentiation and function.

We propose a model whereby Sema3fb signals to Plxna3/Nrp2b-expressing ventricular cardiomyocytes in a cell autonomous manner. In support, *sema3fb* mRNA is expressed throughout the early zebrafish myocardium, while mRNAs for Plxna3 and Nrp2 appear restricted to the ventricle. The timing of when the heart phenotypes present in the *sema3fb* mutants argues against the causative involvement of non-cardiomyocyte tissues. In mouse, Purkinje fibers secrete SEMA3A that patterns sympathetic innervation of the pacemaker node^21^. Sema3fb in zebrafish likely does not play a similar role, in that the cardiomyocyte phenotypes arise prior to when sympathetic nerves innervate the node^69^. Epicardial cells also arrive in the heart after chamber defects are evident in the 48 hpf mutants; epicardial cells emerge at around 55 hpf and migrate to and attach to the myocardium by 72 hpf^33^. Interestingly, Sema3fb is important in zebrafish epicardial development at later stages: by 5 dpf *sema3fb* is expressed by *tbx18*+ epicardial cells and not cardiomyocytes, and Sema3fb regulates epicardial cell numbers in the bulbus arteriosus^70^. We also saw in mutant hearts the normal arrival of cardiomyocytes from non-heart sources, including from the secondary heart field^64^ and CNC^71^. One last possibility is the endocardium. For instance, PLXND1 signaling within the murine endocardium promotes cardiomyocytes to form trabeculae^72^. Endocardial development is delayed initially by the loss of Sema3fb, however, our data argue that alterations in cardiomyocytes do not occur secondary to defects in the adjacent endocardial cells. First, in both WT and *sema3fb* mutants the endocardium lines both heart chambers, and the endocardium-derived heart valves form. The mutant valves do appear mis-distributed at the atrioventricular border, possibly as a result of altered gene expression at the atrioventricular border, but this is a mild phenotype. Second, the cardiomyocytes and not the endocardium express mRNAs for the known Sema3f receptors, Plxna3 and Nrp2^73^. Sema3fb is unlikely to act on endocardial cells via Vegfr2, despite a known inhibitory effect of SEMA3F on VEGF signaling^74, 75^. SEMA3F inhibition of VEGF signaling occurs via NRP and not VEGFR^74, 75^, and the endocardium does not appear to express *nrps*^67^. Finally, we saw no obvious defects in heart tube assembly, a process known to be influenced by the endocardium^76^. These data argue that the contribution of the endocardium to the initial heart impairments we observe in *sema3fb* mutants is minimal. Instead, the literature and our data support strongly the idea that the cardiomyocyte defects arise from the loss of Sema3fb signaling within the cardiomyocytes themselves. Nonetheless, it is possible that defects in valve anatomy and gene expression contribute to the cardiac dysfunction and edema seen in the *sema3fb* mutants.

Our data argue against a Sema3fb signal promoting directly a ventricular fate or repressing atrial/AVC fates. While both *sema3fb* and receptor mRNAs are expressed within the early bilateral heart field, ventricular and atrial cardiomyocyte progenitors are specified much earlier, prior to zebrafish gastrulation^77^. Indeed, both the formation of the ventricular and atrial chambers and their subsequent morphogenesis appear to occurs normally in the absence of Sema3fb. Furthermore, *irx1a* and *tbx5a*, transcription factors implicated in ventricular cardiomyocyte differentiation, remain expressed throughout the mutant ventricle. Thus, if Sema3fb has an early role in heart development, it likely acts redundantly with other mechanisms.

The main impact of Sema3fb loss occurs after the initial specification of the chambers. Both atrial and ventricular chambers are smaller in the absence of Sema3fb, and several genes and proteins whose expression normally obeys the border between the atrium and ventricle are disrupted: 1) *bmp4a* expression fails to refine to the atrioventricular band, 2) *myh7* is largely absent from the ventricular side of the border, 3) *myh6* spreads anteriorly past the atrioventricular constriction, and 4) DM-GRASP and MF20 that are normally expressed preferentially in the ventricle show significant expression throughout the atrium. These data support the idea that Sema3fb provides a patterned signal to ventricular cardiomyocytes at the border with the atrial chamber. The latter lacks Plxna3, and Sema3fb signaling should not be present. Likely the signal provided by Sema3fb does not establish the border between the chambers, as the AVC does form in the *sema3fb^ca305^* heart. Further, several genes still exhibit the expected profound changes in expression at the border: The posterior border of the *irx1a* expression domain obeys the boundary, *bmp4a* mRNA is located preferentially at the AVC, and *myh6* mRNA is largely restricted to the atrial chamber. Instead, we propose that Sema3fb allows for the sharpness of the border to emerge in a timely manner.

Ventricular cardiomyocytes in the border region can respond to Sema3fb while immediately adjacent atrial cardiomyocytes cannot. Our data suggest that this differential signaling ability is required for the refinement of the AVC region. In the absence of border refinement when the Sema3fb signal is absent, the AVC region forms, as exhibited by preferential *bmp4a* expression, but the domain’s border expands into the ventricle and/or the atrium. The Sema3f overexpression data argues against the possibility that Sema3f directly regulates gene expression, as ectopic *bmp4a* expression in the 48 hpf heart is seen with both loss- and gain-of function of Sema3f. Rather, the overexpression data support the idea that Sema3f tells ventricular cardiomyocytes that they sit at the border with the atrium. If the correct spatial information is lost, either because the signal itself is gone (mutant), or the signal is abnormal with cells in the AVC region experiencing different levels of signaling (overexpression), proper refinement of the AVC region fails to occur. An imprecise AVC region leads to disrupted regulation of cardiomyocyte-specific genes in a chamber-defined manner: DM-GRASP, *myh7*, *myh6* and MF20. In support, restriction of Bmp signaling to the AVC is necessary to prevent inhibition of cardiomyocyte differentiation by Tbx2 within the ventricular chamber^78–80^.

Interestingly, in *sema3fb* mutants the atrium is also impacted with regard to size, functional parameters, and misexpression of ventricular and AVC markers. Given that atrial cardiomyocytes lack the machinery to respond to Sema3fb, altered atrial differentiation arises presumably secondarily to disrupted ventricular cardiomyocyte development. One possibility is that disrupted ventricle function impacts atrial cardiomyocytes, as the function of the two chambers shows some inter-dependence^49^. Explanted mutant atria are still smaller than their WT counterparts, however, despite having removed flow as a variable. Potentially, alterations in the secretion of factors from ventricular cardiomyocytes and/or cells of the AVC affects the differentiation of the nearby atrial cardiomyocytes. In support, both MF20 and DM-GRASP proteins appear upregulated in the atrium of *sema3fb* mutants.

The heart chambers are smaller in *sema3fb* mutants, though the data suggests that fewer cardiomyocytes is not the explanation. The expression of *fgf8a* and *tbx5a* (data not shown) in the primary heart field, and markers of secondary heart field-derived cells, are comparable in the two genotypes. Further, cell death and proliferation at 48 hpf are not impacted by Sema3fb loss. Indeed, similar numbers of ventricular cardiomyocytes are present in WT and mutant hearts at this stage. Instead, without Sema3fb these cardiomyocytes appear to be smaller than normal. The fact that the atrial chamber was smaller in the absence of changes in proliferation, apoptosis and cell integration also argues for smaller cardiomyocytes within the atria. Heart dysfunction could result in smaller cells, in that zebrafish cardiomyocytes hypertrophy normally at about 45 hpf in response to shear stresses arising from the onset of blood flow through the heart^61^. Yet, the atria of explanted mutant hearts were still smaller than those of WT, even though shear force was removed as a functional regulator. Potentially, Sema3fb signaling causes cytoskeletal rearrangements that induce ventricular cardiomyocyte expansion, but our data argue that atrial cardiomyocytes are not responding directly to Sema3fb. For atrial cardiomyocytes, and possibly their ventricular counterparts, disrupted expression of differentiation genes remains a potential explanation for the failure of cardiomyocytes to undergo hypertrophy.

The multigenic and heterogenous disease Hypoplastic Left Heart Syndrome (HLHS) is characterized by a hypoplastic left ventricle, aorta and mitral valve, and without treatment, is generally fatal^81, 82^. The Ohia mouse harbours mutations in *SAP130* and *PCDH9A*, and presents with hypoplasia of the left ventricle, likely due to an intrinsic deficit in cardiomyocyte proliferation and differentiation^83^. Our data support the idea of defective differentiation of ventricular cardiomyocytes as a potential contributor to HLHS. Different from what is often observed in HLHS, however, the ventricular defect in the *sema3fb* mutants translates to a smaller atrium. This involvement of the atrium may reflect a critical interdependence of the atrium and ventricle in the two-chambered zebrafish heart. While *sema3fb* mRNA in the ventricle is largely downregulated by 72 hpf (data not shown), earlier defects in differentiation generates ventricular cardiomyocytes that fail to maintain normal cardiac function as the fish mature. The fact that the edema in these fish appears to resolve, and the mutant alleles are viable, argues for mechanisms that ultimately compensate for the loss of Sema3fb.

Here we provide evidence for a secreted Sema functioning as an extrinsic regulator of regional differentiation in the cardiovascular system. Sema3s are known to be important for development of the mouse cardiovascular system^16^. In some cases, the roles are myocardium-independent, as is the case for SEMA3A and innervation of the heart^21^, and Sema3d and SEMA3C in CNC-dependent generation of cardiomyocytes^84, 85^. Here, we suggest a novel cardiomyocyte autonomous function of Sema3fb in governing the molecular differentiation of cardiomyocytes. While *Sema3f* is expressed in the cardiac cushions and in scattered cells throughout the myocardium of the embryonic chick heart^86^, no cardiac phenotype is described in the Sema3f mouse mutant^87^, nor is Nrp2 expressed by murine cardiac muscle^88^. In mouse, the transmembrane SEMA6D promotes cardiomyocyte proliferation^89^. In concert with our findings, these data support the idea that Semas directly regulate cardiomyocyte development. Additional roles for Semas will likely be identified in heart development as more subtle cardiovascular defects are assessed in loss-of-function models.

## Acknowledgements

We would like to thank Drs. G. Bertolesi, S. Childs and K. Atkinson-Leadbeater for helpful comments on the manuscript, to Dr. S. Childs and Bertolesi for genetic analysis of the *sema3fb* mutant alleles, and to Dr. S. Childs for use of her fish room and confocal microscope. We would also like to thank F. Alshemali and Dr. R. Rose for training on and access to their Leica DM5500 B microscope. The work was supported by a Canadian Institutes of Health Research project grant to S.M., and salary awards from Alberta Innovates-Health Solutions and the Hotchkiss Brain Institute T. Chen Fong Studentship Award to R.H, and from the Natural Sciences and Engineering Research Council of Canada to P.B.C.

---

**Supplemental Figure 1:**
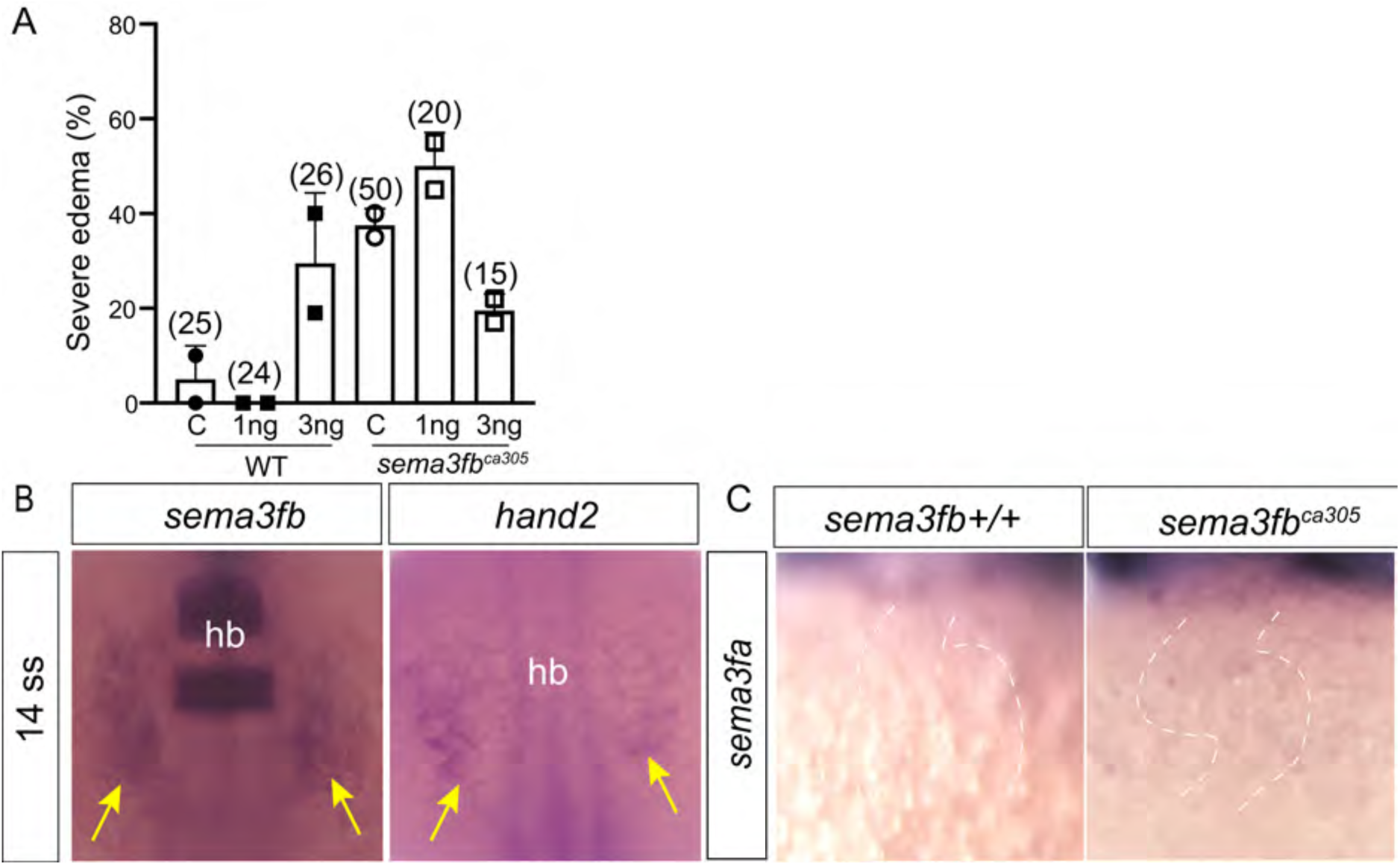
*sema3f* expression and function in the developing heart. **A)** Percentage of embryos with a severe heart edema at 48 hpf in WT and *sema3fb^ca305^* embryos and those injected with 1 ng/ml or 3 ng/ml of an antisense morpholino oligonucleotide against the ATG start site of the *sema3fb* gene. Numbers in brackets represent number of embryos (N=2 independent experiments). **B)** Dorsal view of whole mount ISH for *sema3fb* at the 14 somite stage reveals that mRNA is expressed in the early heart field (yellow arrows) as marked by *hand2*. **C)** *sema3fa* shows little or no expression in the 48 hpf WT and *sema3fb* mutant heart.

**Supplementary Figure 2:**
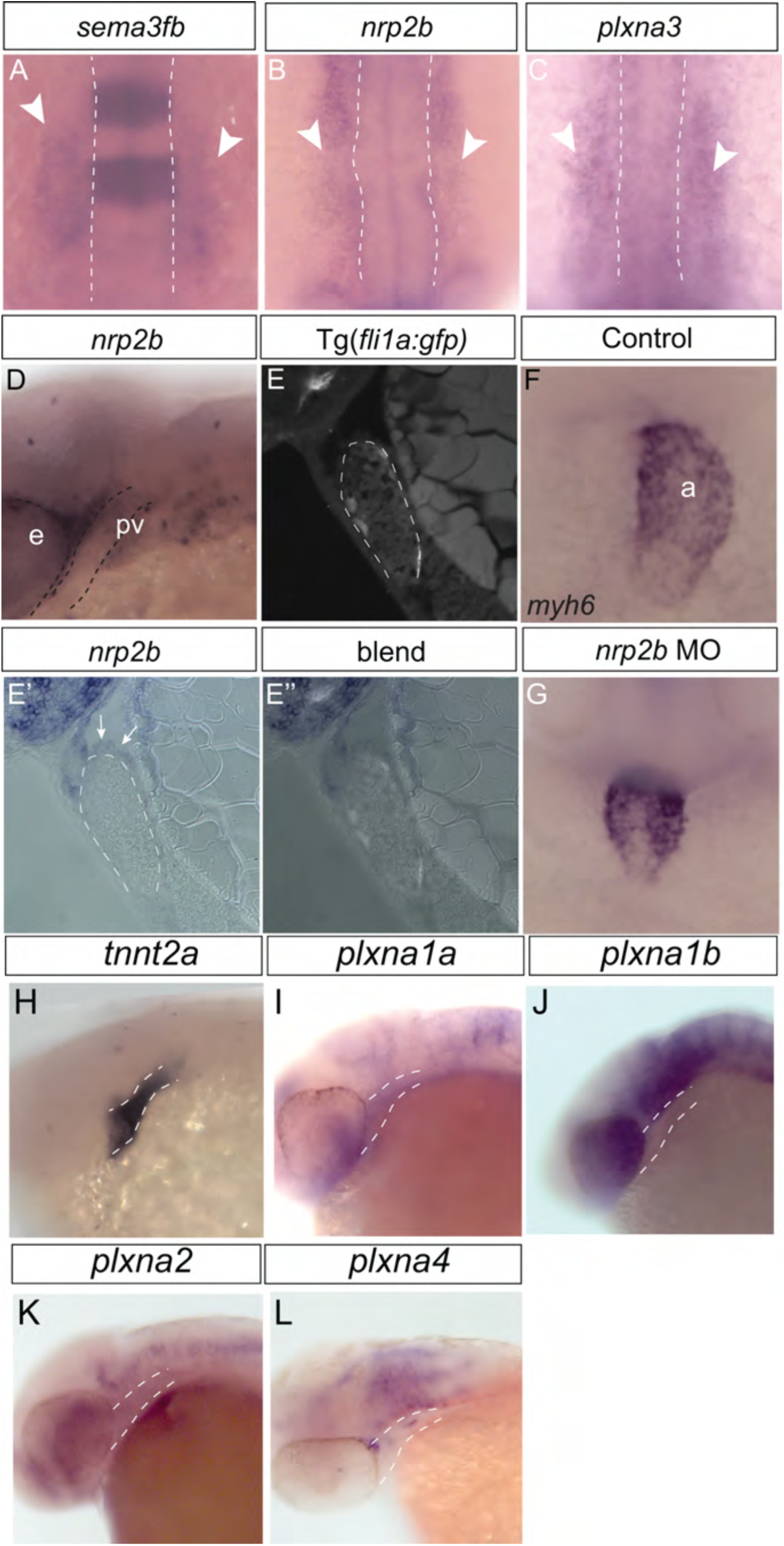
Early heart development occurs normally in the absence of Sema3fb. Dorsal (A-C) and lateral (D-F) views of 13 hpf WT (A,D), *sema3fb^ca305^* (B, E), and *sema3fb^306^* (C, F) embryos processed by whole mount ISH. **A-F)** *fgf8a* is expressed normally (N=2, n=10) in the heart field containing anterior lateral plate mesoderm as indicated by arrowheads in low magnification lateral (D-F) and high magnification dorsal (A-C) views. A, anterior; mhb, midbrain-hindbrain boundary; som, somites; tb, tailbud. Scale bar: 50 µm for A-C, 100 µm for D-F.

**Supplemental Figure 3.**
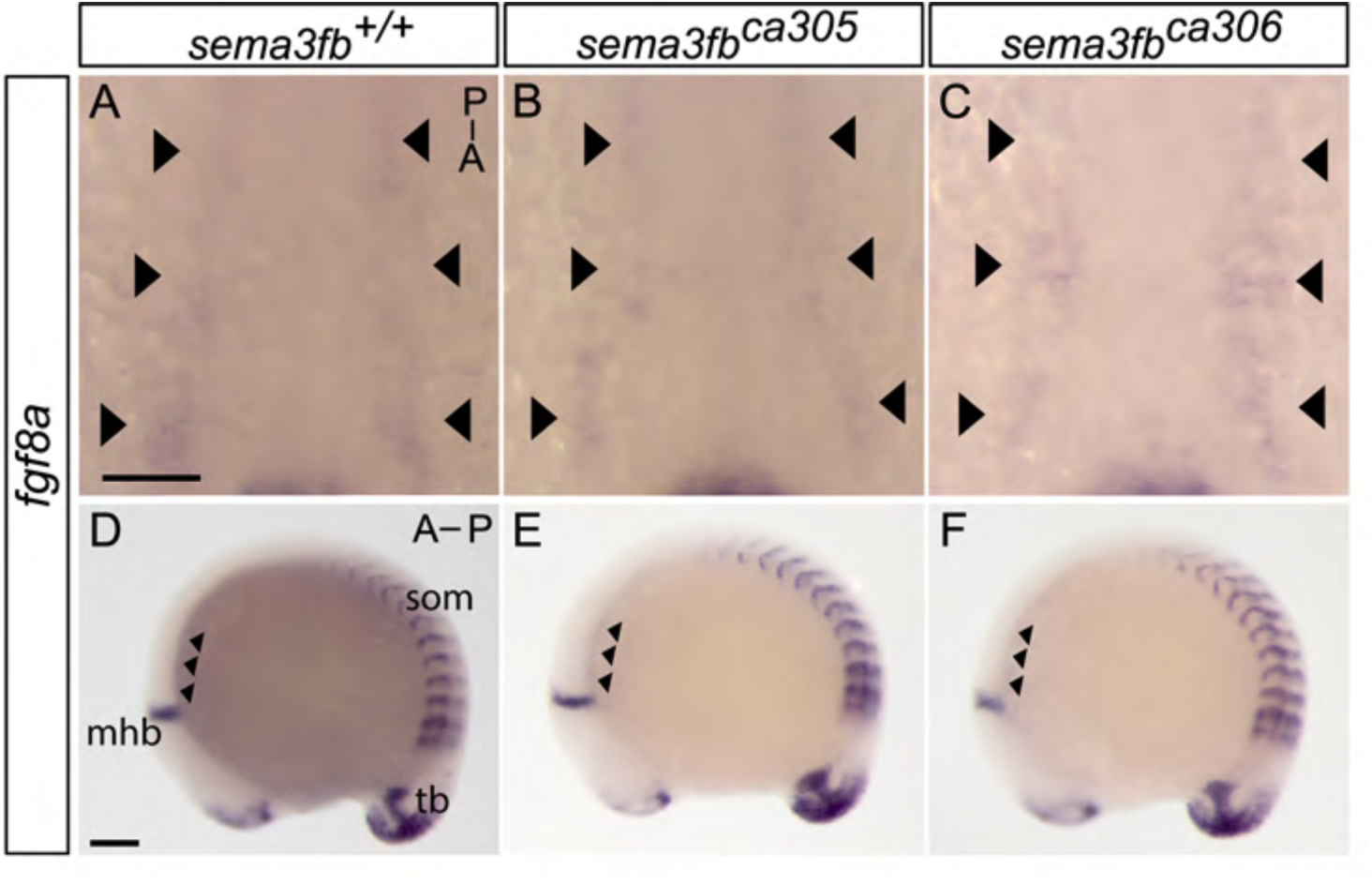
Heart tube extension is delayed slightly in sema3fb mutant embryos. Ventral views of WT, *sema3fb^ca305^* and *sema3fb^ca306^* 24 hpf embryos processed for whole mount ISH using cardiomyocyte specific riboprobes. **A-C)** *tnnt2a* ISH shows that the heart elongates in the WT to form a tube (N=3, n=15/17) (A). Progression into the extension stage is defective at 24 hpf in both *sema3fb* mutant alleles (N=3, 14/21 *sema3fb^ca305^*; N=3, 15/21 *sema3fb^ca306^*), with cone (B) and condensed (C) morphologies being exhibited. **D)** Percent occurrence of the different cardiac morphologies. **E-G)** Ventricular cardiomyocyte specific *myh7* riboprobe also reveals elongation of WT hearts (N=3, n=17/20) (E), but that *sema3fb* mutant hearts exhibit cone or condensed morphologies (N=3, 12/19 *sema3fb^ca305^*; N=3, 13/19 *sema3fb^ca306^*) (F-G). **H)** Percent occurrence of the different cardiac morphologies. **I-K)** Atrial cardiomyocyte specific *myh6* riboprobe reveals that WT atria are tubularized (N=3, n=17/20) (I), but are primarily at the cone stage in both the *sema3fb* mutant alleles (N=3, 19/20 *sema3fb^ca305^*; N=3, 11/18 *sema3fb^ca306^*) (J-K). **L)** Percent occurrence of the different cardiac morphologies. **M-O)** Ventral views of embryos processed for whole mount ISH for the ventricle specific marker *myh7*. Heart tube elongation was scored for WT embryos at 23 hpf (N=1, n=9/10, M), and at 26 hpf for *sema3fb^ca305^* (N=1, n=8/10, N) and *sema3fb^ca306^* (N=1, n=9/10, O) embryos. **P-Q)** Ventral views of DM-GRASP whole mount immunostaining of embryos at 24 hpf show cranial ganglion (red asterisk) that has migrated and formed normally in WT (n=4, P) and *sema3fb^ca305^* (n=4, Q) embryos, despite the heart of mutants showing a mild developmental delay (yellow arrowhead). R) Percentage occurrence of *myh7*-labeled heart morphologies between WT at 23 hpf, and *sema3fb^ca305^* and *sema3fb^ca306^* at 26 hpf. A: anterior; P: posterior. Scale bar: 100 µm.

**Supplemental Figure 4.**
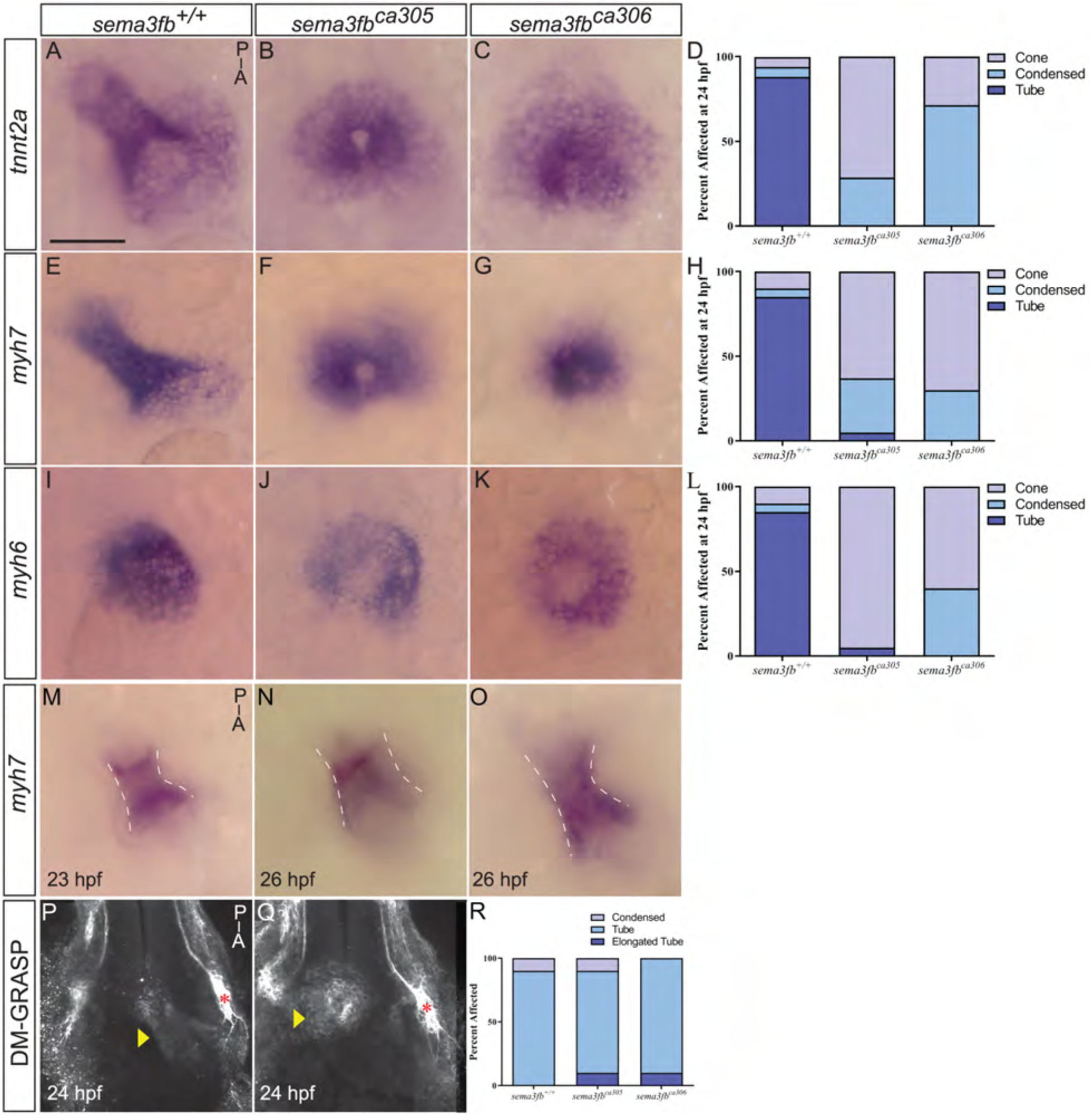
Ventricle size is reduced significantly in *sema3fb* mutants. **A-D)** Ventricle measurements were made from still images taken during video microscopy of WT and mutant hearts during diastole (A,C) and systole (B,D) at 72 hpf. Quantitation reveals that in mutants as compared to WT the ventricle diameter during diastole (A, **p=0.0017) and during systole (B, **p=0.0063) is significantly smaller, and ventricle length is significantly shorter at diastole (C, *p=0.048) and systole (D, *p=0.031). Error bars are SEM over 3 independent experiments (n=12 WT, n=20 *sema3fb^ca305^*). Statistics represent the Mann-Whitney U test.

**Supplemental Figure 5.**
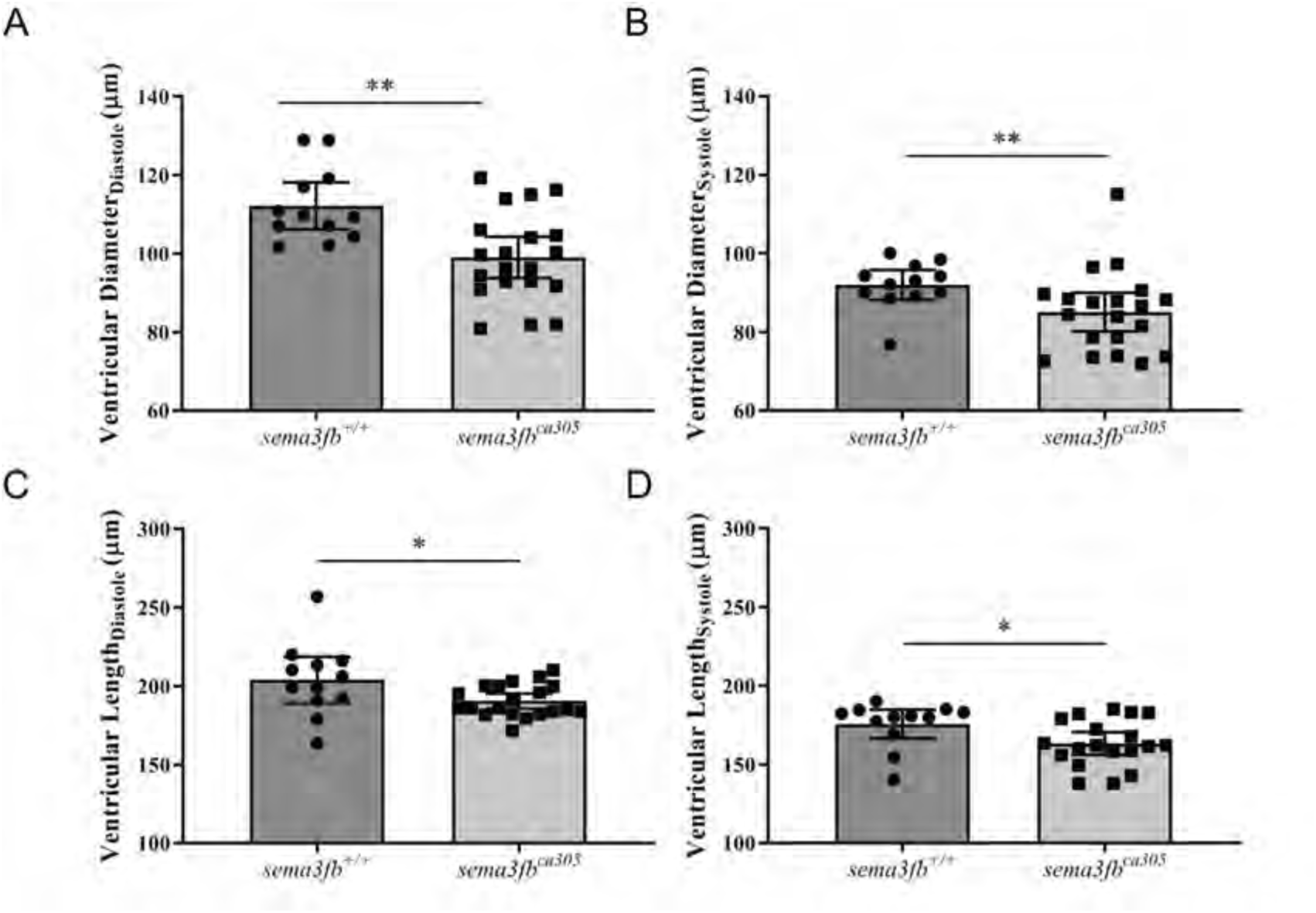
Cardiomyocytes derived from the secondary heart field and cardiac neural crest are present in the developing *sema3fb* mutant heart. Whole mount ISH of 48 hpf hearts with riboprobes against the CNC marker *crip2* (A,B) and the secondary heart field marker *ltbp3a* (C,D). Similar labeling is seen in WT and *sema3fb^ca305^* hearts for *crip2* (A-B, N=1, n=11 WT and N=1, n=11 *sema3fb^ca305^*) and *ltbp3a* (C-D, N=1, n=6 WT and N=1, n=7 *sema3fb^ca305^*). A: anterior; P: posterior. Scale bar: 50 µm.

**Supplemental Figure 6.**
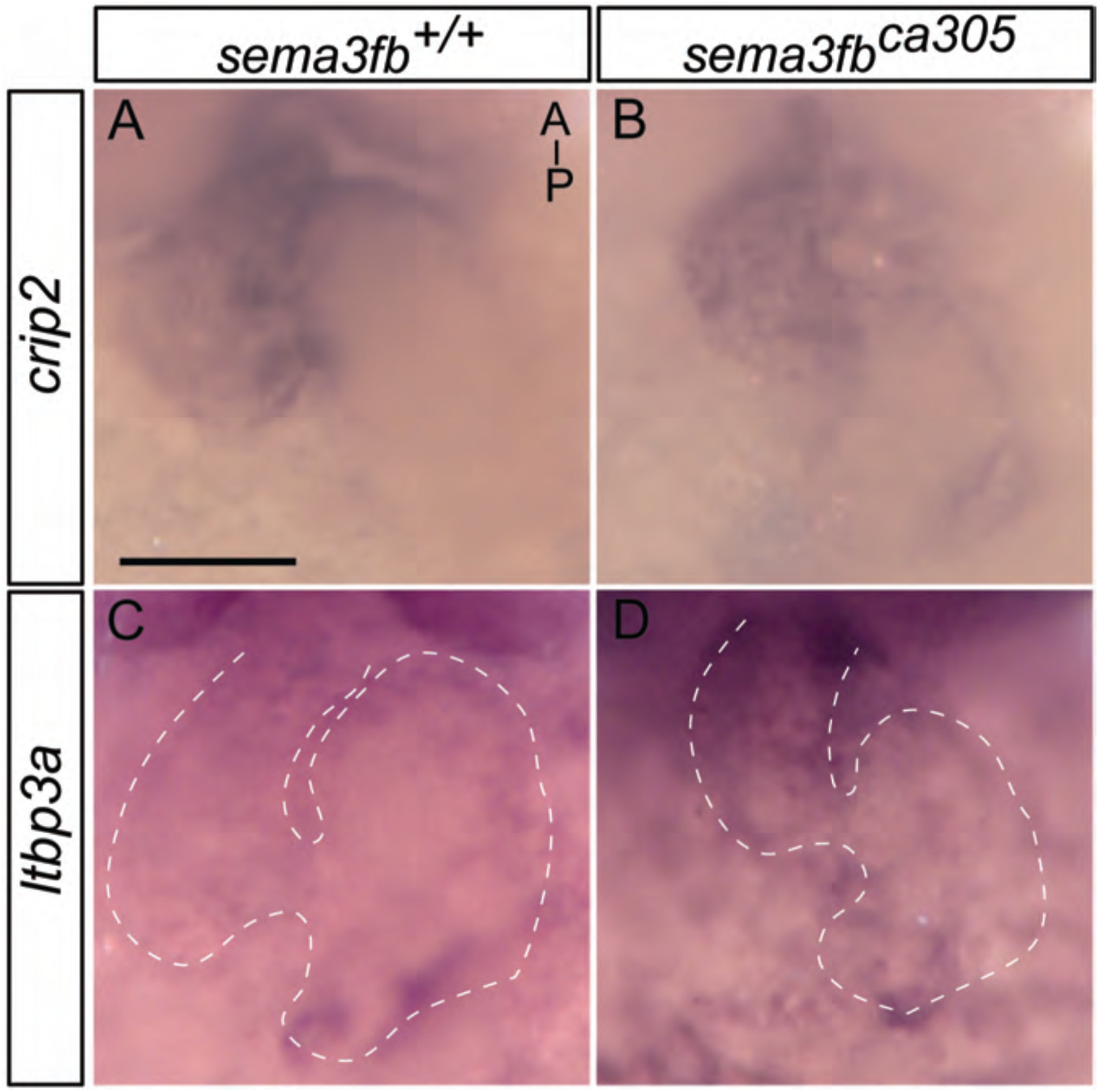
Reduced cardiac chamber size is due to intrinsic loss of Sema3fb. Heart explants dissected from embryos at 24 hpf and cultured for 24 hours before processing by whole mount ISH for the atrial cardiomyocyte marker *myh6*. The atria of *sema3fb^ca305^* hearts (N=2, n=11/13) are smaller in size than WT atria (N=2, n=8/8). (C) Graph of atrial area shows a significant reduction in size in *sema3fb^ca305^* (*p=0.025, Mann-Whitney U test) as compared to WT explants. Error bars represent SEM. Scale bar: 50 µm.

**Supplemental Figure 7:**
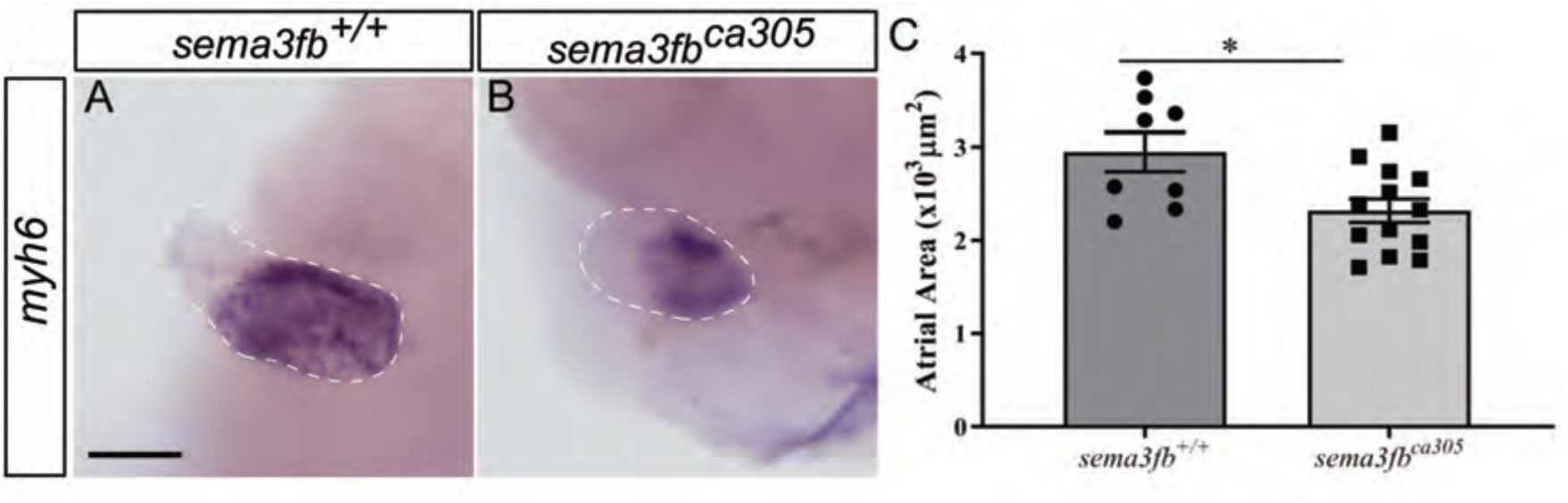
*nrp2b* is expressed by ventricular cardiomyocytes and is required for normal heart development. **A-C)** Dorsal views of whole mount ISH for *sema3fb*, *nrp2b* and *plxna3* at 18 hpf, revealing expression in the developing heart field (white arrowheads). **D-E)** *nrp2b* ISH signal in the presumptive ventricle as viewed laterally in a 28 hpf embryo (D), and in a sagittal section (E’) of a 36 hpf *Tg(flk:EGFP)* embryo where endocardial cells that line the heart and neighbour the cardiomyocytes are GFP positive (E). A blend of the two signals is shown in E’’. **F-G)** *myh6* labelling of the atrium in a control embryo (F) and one injected at the one-cell stage with an antisense MO against *nrp2b* (G). **H-L)** ISH for various *plxna* receptors (I-L) show minimal signal in the *tnnt2a* (H) labeled heart at 24 hpf.

